# A diverse proteome is present and enzymatically active in metabolite extracts

**DOI:** 10.1101/2023.12.22.573071

**Authors:** Rachel (Rae) J. House, Molly T. Soper-Hopper, Michael P. Vincent, Abigail E. Ellis, Colt D. Capan, Zachary B. Madaj, Emily Wolfrum, Christine N. Isaguirre, Carlos D. Castello, Amy B. Johnson, Martha L. Escobar Galvis, Kelsey S. Williams, Hyoungjoo Lee, Ryan D. Sheldon

## Abstract

Metabolomics, a foundational tool in metabolism research, relies on the accurate transmittal of biochemical profiles underlying biological phenotypes. Over the years, workflows used in metabolomics have been assumed to remove enzymes to preserve metabolite levels during processing. Here, we uncover a diverse landscape of over 1,000 proteins, strongly enriched for metabolic enzymes, within metabolite extracts generated using common extraction workflows. Moreover, by combining in-extract stable isotope additions and enzyme inhibitors, we demonstrate transaminase activity, which is preventable by protein removal by 3 kDa filtration. We extend these findings to untargeted metabolomics, revealing that both post-extraction formation of glutamate dipeptide and depletion of total glutathione can also be prevented by removing proteins. Finally, we present a simple yet novel workflow that integrates passive filtration for protein removal of crude metabolite extracts as a superior method for broad-coverage metabolomics. Our findings have broad-reaching experimental implications across all fields that use metabolomics and molecular metabolism, especially cancer, immunology, and diabetes research.

## Introduction

Metabolism is a highly dynamic, interconnected, and diverse set of chemical reactions required for energetic homeostasis, redox balance, macromolecular anabolism, and cellular signal transduction. Metabolic phenotypes are not only components but also key drivers of physiological and pathological states. Interrogation and experimental manipulation of metabolism has enhanced our understanding of topics including exercise [1], cancer [2–7], and immune function [8–10], and has paved the way for metabolism-based therapeutics [11]. Reliable and reproducible detection of biologically relevant metabolic phenotypes is central to the continued success of metabolism research.

Mass spectrometry-based metabolomics enables detection and quantification of hundreds to thousands of metabolites in a sample. Unlike other ‘omics’ techniques, no single workflow captures the entire metabolome due to its vast chemical diversity. Consequently, countless metabolomics workflows have been developed to measure different slices of the metabolome. Across workflows, metabolomic phenotyping is predicated on the preservation of biological phenotype throughout quenching, extraction, and chromatography-mass spectrometry analysis. Therefore, it is essential to evaluate metabolomics workflows based on their preservation of biological phenotype.

In a metabolomics workflow, the first step involves quenching metabolism, commonly by freezing. After metabolism is quenched, compounds of interest are extracted from the biological matrix. The roots of contemporary metabolite extraction can be traced back to the early 1900s, when pioneers like Schonheimer developed innovative methods for metabolic physiology and stable isotope tracing analyses [12–15]. Today, in the era of high-resolution mass spectrometry and untargeted metabolomics, the experimental prerogative is to maximize compound coverage while minimizing orthogonal workflows. This is a challenging goal given (i) metabolite chemical diversity (hydrophobicity, polarity, size, etc.), (ii) metabolite chemical similarity, which can result in spontaneous or process-induced metabolite interconversions, and (iii) the need for compatibility between extraction modality and downstream analytical workflows. As a result, available extraction workflows prioritize everything from recovery of a specific compound to global metabolome coverage [16–25].

Of the published extraction modalities, the most common are monophasic polar solvent extractions (e.g., acetonitrile and/or methanol) [26, 27] and biphasic liquid-liquid extractions (e.g., chloroform, methanol, and water) [28]. However, even within polar and biphasic extraction methods, the procedural details vary wildly, including water content, additives (acid, base, ETDA, etc.), solvent ratio, sample to solvent ratio, and pH. Given the wide range of metabolite extraction methodologies, is essential to understand the effect of extraction diversity on metabolomic analytical scope, reproducibility, and biological insight.

Despite disparate procedural details, all metabolite extraction methods include a step to precipitate proteins and other unwanted biomolecules, such as RNA, DNA, glycogen, and lipids. This critical experimental step is generally assumed to be complete. Indeed, while many studies demonstrate that the insoluble extract fraction contains protein and macromolecules, no study has provided empirical evidence to support the assumption that the soluble, metabolite-containing fraction is devoid of *active* proteins [29–36]. A single report demonstrated that 2-6% of total serum proteins remain in the soluble metabolite fraction in an extraction solvent-dependent manner [37]. If protein carryover were to occur in other matrices with metabolic enzyme-enriched proteomes (e.g., cells or tissues), it is plausible that metabolomics extracts are, in fact, unintentional bioreactors of enzymes and their substrates.

Here, we uncover a diverse proteome in metabolite extracts across a wide range of sample and extraction types. Strikingly, these metabolite extract proteomes are enriched for conditionally active metabolic enzymes that can cause in-extract metabolite interconversions. We also demonstrate that post-extraction metabolite interconversions are preventable by removing proteins using 3 kDa filtration. Finally, we present a novel method for broad-coverage metabolomics that uses passive filtration to remove proteins from metabolite extracts.

## Results

### Metabolomic responses to extraction water content are not compound-intrinsic

Metabolite extraction constrains the analytical scope of metabolomics experiments, and optimization of extraction conditions may enhance metabolome coverage. To this end, we sought to evaluate how extraction conditions would affect detection of polar metabolites, such as nucleotides. We hypothesized that increased extraction water content during extraction would improve recovery these compounds. Using cryopulverized and pooled mouse liver tissue as a model system, we extracted metabolites with a widely used extraction approach of 40% acetonitrile, 40% methanol, and 20% water (AMW20) [38–50]. Then, we performed a secondary addition of water to the crude extract immediately after homogenization to achieve final water content between 25% (AMW25) and 60% (AMW60) in 5% increments (**Figure 1A**). Control experiments with addition of a second volume of AMW20 solvent confirmed that metabolomic changes were not due to increased extraction volume alone (**Figure S1**).

**Fig. 1.**
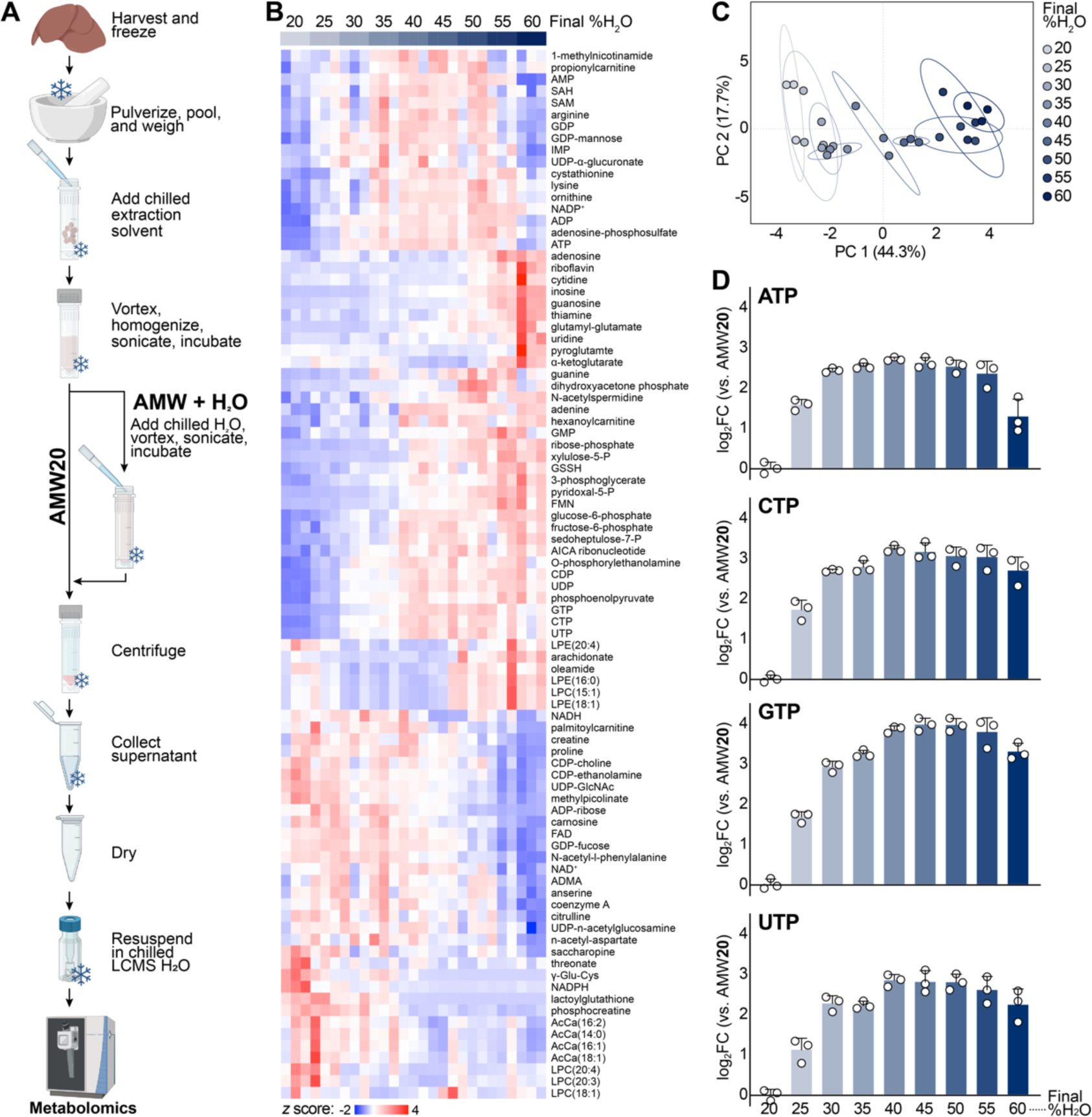
Extraction water content strongly influences metabolomics. (**A**) Schematic of AMW and AMW + H_2_O extractions. (**B**) Heatmap depicting relative abundance of murine liver metabolites across extraction conditions, from AMW20 to AMW60. All significantly different metabolites are shown. Significance was calculated by ANOVA using MetaboAnalyst 5.0, and row hierarchical clustering was performed using Morpheus (Broad). (**C**) PCA of AMW20-AMW60 metabolites in murine liver (95% confidence ellipse, *n*=3 per group). (**D**) Relative abundance of murine liver nucleotides across extraction conditions, from AMW20 to AMW60 (*versus* AMW20, mean ± SD, *n*=3 per group).

Extraction water content significantly affected peak areas of 89 of 195 metabolites detected (**Figure 1B**) and led to stratification in the first principle component (44.3%) by PCA (**Figure 1C**). We specifically noted a water dose-dependent increase in nucleotide triphosphates, with the greatest stepwise increase occurring between AMW20 and AMW30 (**Figure 1D**). This large increase (5- to 8-fold) in nucleotide triphosphate signal over a relatively small range of extraction water content (20-30%) highlights the dependency of metabolomics experiments on extraction conditions and the need to tightly control such variables.

Given the complex metabolomic effect of water titration (**Figure 1**), we hypothesized that compound hydrophobicity dictates the metabolome responsiveness to extraction water content. To test this hypothesis, we evaluated the change in metabolite response to water (AMW50/AMW20) as a function of LogP, the partitioning coefficient between octanol (positive LogP, hydrophobic) and water (negative LogP, hydrophilic). Indeed, nucleotides increase in abundance as a function of extraction water content, and this is dependent on phosphate group number (i.e., nTP>nDP>nMP) (**Figure 2A**). Additionally, acyl-carnitines become increasingly hydrophobic with acyl-chain length and their detected abundance decreased with extraction water content (**Figure 2B**). To explore this further, we used a cheminformatics approach to broadly examine the relationship between metabolite hydrophobicity and water content-mediated changes in LCMS-measured abundance (**Figure 2C**). The compound SMILES (Simplified Molecular Input Line Entry System) was used to predict the LogP values, which ranged from −4.9 (N-Glycolylneuraminic acid; most hydrophilic) to

**Fig. 2.**
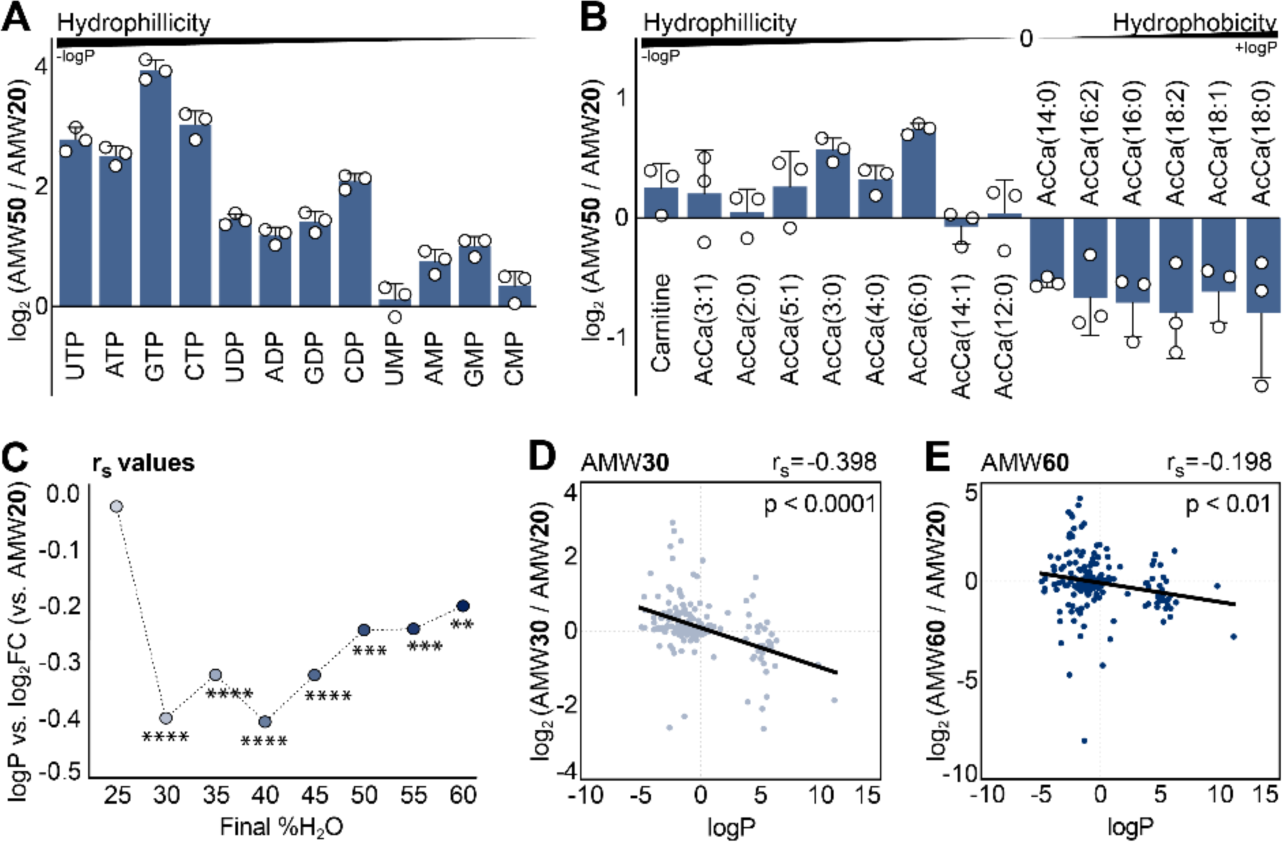
Compound non-intrinsic factors influence metabolite response to extraction water content. (**A**) Relative abundance of AMW50 nucleotides (*versus* AMW20). Nucleotides are ordered by PubChem XlogP3 value (mean ± SD, *n*=3 per group). (**B**) Relative abundance of AMW50 carnitine and acyl-carnitine species (*versus* AMW20). Carnitine and acyl-carnitine species are ordered by ChemAxon LogP value (mean ± SD, *n*=3 per group). (**C**) Metabolite hydrophobicity and recovery correlation in AMW25-AMW60 extraction conditions. Spearman correlation between the LogP and the specified mean Log2 fold change (X = 4:4:2 + X% water *versus* the 4:4:2 +20% water control) as a function of water content. ****p < 0.0001; ***p < 0.0005; **p < 0.01. (**D**) Mean Log2 fold change (*n*=3) of murine liver metabolites in the AMW30 extraction is shown with respect to the control condition (AMW20). Spearman correlation coefficient (r_s_), *p*-value, and linear regression model (black line) are displayed. (**E**) Mean Log2 fold change (*n*=3) of murine liver metabolites in the AMW60 extraction is shown with respect to the control condition (AMW20). Spearman correlation coefficient (r_s_), *p*-value, and linear regression model (black line) are displayed.

11.2 (phosphatidyl-choline, PC[16:0_18:1]; most hydrophobic) (**Figure S2**). We hypothesized an inverse relationship between the LogP and the Log2FC with respect to AMW20, which would support the idea that recovery/enrichment of more hydrophilic compounds would increase with water content whereas the recovery of more hydrophobic compounds (higher LogP values) would decrease with water content. Consistent with this, a significant inverse correlation was observed beginning at AMW30 (**Figure S3**). However, instead of becoming more negative, this correlation weakened with increasing water content (**Figure 2D; Figure S3**). This trend is surprising and suggests that non-compound intrinsic factors, arising from increased water content, influence compound recovery during extraction.

Another surprising observation from these studies was the extraction water-dependent loss of the internal standard, D_5_-Glutamate, and concomitant gain of D_4_-Glutamate (**Figure 3A**). Unlabeled glutamate from the sample was unaffected (**Figure 3B**), indicating that glutamate is stable and unaffected by extraction water content in this range. As further controls, another deuterated internal standard, D_8_-tryptophan, was unchanged (**Figure 3C**) and extract pH was unaffected by water content (**Figure S4**), collectively suggesting that D_5_ → D_4_-glutamate transition was not due to spontaneous hydrogen-deuterium exchange. Given our observations that non-compound intrinsic factors contribute to extraction water responses (**Figure 2**) and water-dependent conversion that D_5_ → D_4_-glutamate, we next considered whether a protein mediated mechanism could be responsible.

**Fig. 3.**
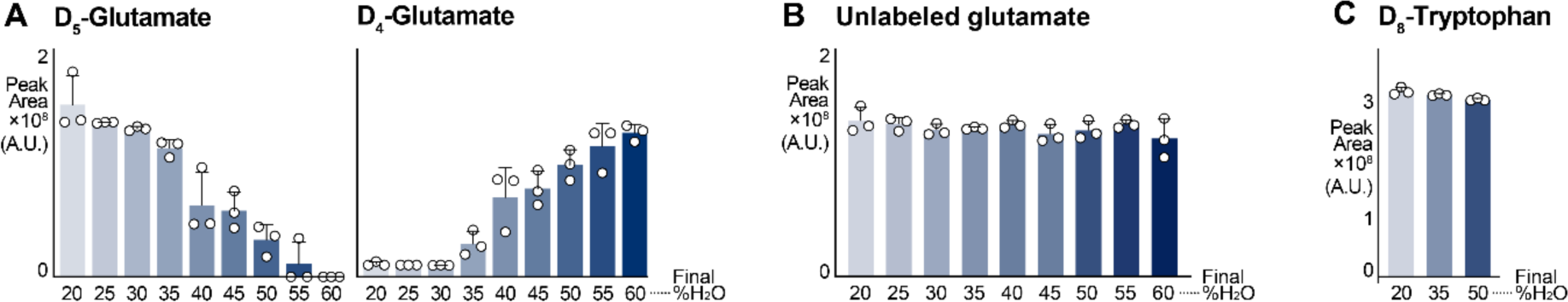
Single deuterium loss from D_5_ glutamate is extraction water dose dependent. (**A**) Relative abundance of D_5_-glutamate and D_4_-glutamate across extraction conditions, from AMW20 to AMW60. D_5_-glutamate was added at resuspension (mean ± SD, *n*=3 per group). (**B**) Relative abundance of unlabeled, sample originating glutamate across extraction conditions, from AMW20 to AMW60. (mean ± SD, *n*=3 per group). (**C**) D_8_-tryptophan is unaffected by extraction water content (mean ± SD, *n*=3 per group).

### Proteins remain in the metabolite fraction during extraction

It is generally assumed that protein precipitation during metabolite extraction is complete or at least is inconsequential. To determine if the observations in **Figure 2** were in fact due to enzymatic activity, we assessed the total protein content of AMW20 and AMW50 extracts (**Figure 4A**). This revealed 11.3 µg protein/mg liver and 15.7 µg/mg of liver, respectively, compared to 153.8 µg/mg of whole liver lysates (**Figure 4B**). Since the BCA reagent also reacts with non-proteinaceous peptides such as glutathione, which is abundant in the liver [51], we filtered crude extracts through a 3 kDa filter to remove proteins, resulting in a 20-30% decrease in calculated BCA protein content (**Figure 4A-B**). To understand the fraction of BCA signal in AMW extracts arising from proteins, we subtracted the 3 kDa eluate BCA signal from total extract BCA signal and found that 2.7 and 4.1 µg of >3 kDa proteins per mg of liver tissue are present in AMW20 and AMW50 extracts, respectively (**Figure 4C**).

**Fig. 4.**
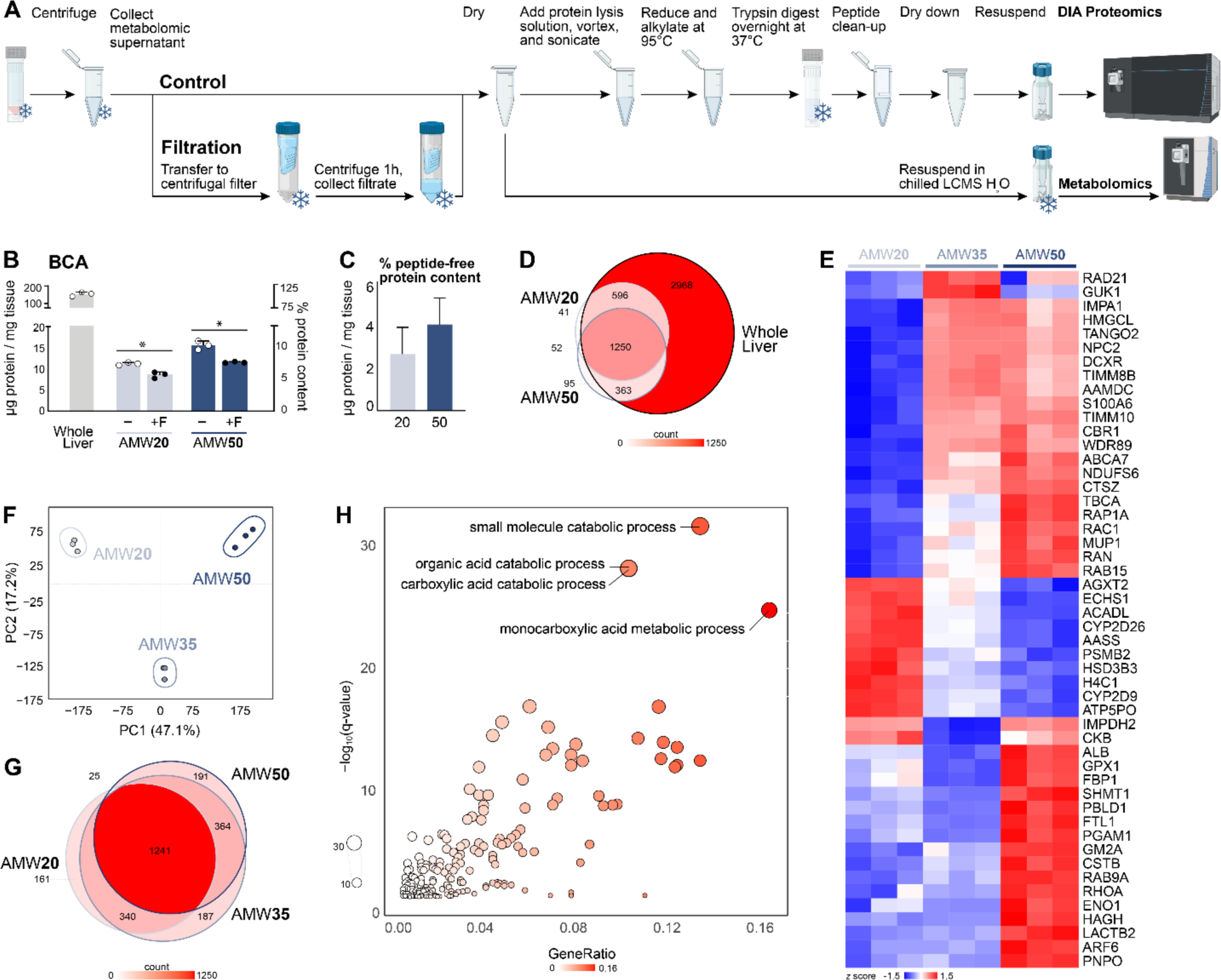
Proteomic content of metabolite extracts. (**A**) Schematic of sample preparation post-metabolite extraction for sample filtration, and bottom-up proteomics analysis. (**B**) BCA determination of protein content normalized to starting tissue amount. Significance calculated by Welch’s t-test. Percentage protein content calculated as a fraction of the whole liver extract protein content; *n*=3 per group. (**C**) Peptide free protein content in AMW20 and AMW50 metabolite extractions; calculated as the average mg protein/mg tissue of the unfiltered extraction minus the average filtered extraction and normalized to the average total protein content in a whole liver (*n*=3 per group). (**D**) Number of protein groups identified in AMW20, AMW50, and whole liver extracts; *n*=3 per group. (**E**) Heatmap depicting relative abundance of murine protein groups quantified across extraction conditions and row standardized. The top 50 most differentially abundant proteins (based FDR p-value) are shown. Significance calculated via LIMMA-eBayes on Log2 transformed proteins without any missing values. (**F**) PCA of AMW20, AMW30, and AMW50 protein groups in murine liver (95% confidence ellipse, *n*=3 per group). (**G**) Number of protein groups identified in AMW20, AMW35, and AMW50 metabolite extracts (*n*=3 per group). (**H**) Gene set enrichment analysis (biological processes) of proteins significantly impacted by water content. The top 5 pathways (based on FDR q-values) are labelled. Points are sized by –Log10(q-value) and colored by the ratio of differentially abundant proteins divided by total proteins in each pathway (GeneRatio).

Next, we used quantitative proteomics to evaluate the protein composition of AMW20 and AMW50 extracts. We detected 1,939 and 1,760 proteins in AMW20 and AMW50 extracts, respectively, compared to 5,177 in whole liver (**Figure 4D**). 3 kDa filtration of extracts confirmed the removal of most of these proteins (**Figure S5**). To better understand the relationship between the extraction water content and the composition of the protein extracts, we added an intermediate AMW extract with 35% final water content (AMW35) and performed proteomics on the extracts. Differential abundance analysis of quantified protein abundances across the three extraction types revealed that proteins were highly responsive to water content, with 1,084 significantly affected (**Figure 4E-F**). Further, while the majority (1,241) of proteins were detected in all three extraction modalities, 161, 187 and 191 proteins were unique to AMW20, AMW35, and AMW50 extracts (**Figure 4G**). To gain clarity on the types of proteins present in metabolite extracts we performed gene set enrichment analysis (GSEA). Strikingly, the top pathway was “small molecule metabolic process”, and each of the top ten pathways were metabolic (**Figure 4H**).

### Enzymatic activity occurs post-extraction

We noted the presence of eleven transaminases among the proteins found in metabolite extracts (**Figure 4**). This included the aspartate-glutamate transaminase (GOT1), which was elevated in AMW50 *versus* other conditions (**Figure 5A**). Transaminases are a class of enzymes that transfer the amine nitrogen of an amino acid to a corresponding ketoacid (e.g., glutamate → α-ketoglutarate) using pyridoxal-5-phosphate (PLP) as a cofactor. Interestingly, we noted similar water-dependent increases in PLP content (**Figure 5B**). The concomitant elevation of enzyme and cofactor in AMW50 led us to hypothesize that in-extract transaminase activity could be responsible for the D_5_ → D_4_-glutamate transition observed in **Figure 3**. Indeed, D_5_-glutamate contains five deuterated hydrogens, including one on the α-amino carbon that would be lost through deamination (**Figure 5C**). To test this hypothesis, we monitored the loss of D_5_-glutamate and appearance of D_4_-αKG and D_4_-glutamate using high-resolution LCMS. Consistent with glutamate transamination, we noted a loss in D_5_-glutamate abundance ([M-H] = 151.0774 m/z), and a concomitant gain in D_4_-glutamate ([M-H] = 150.0711 m/z) in AMW50 *versus* AMW20 (**Figure 5D**). Finally, to causally link this D_5_ → D_4_-glutamate shift to sample protein content and specifically to transaminase activity, we repeated this experiment in AMW50 extracts that were either 3 kDa-filtered or in which the pan-transaminase inhibitor aminooxyacetic acid (AOA) [52] was added. Remarkably, both protein removal and transaminase inhibition independently preserved D_5_-glutamate and mitigated the appearance of D_4_-αKG and D_4_-glutamate at 24 hr post-resuspension (**Figure 5E**).

**Fig. 5.**
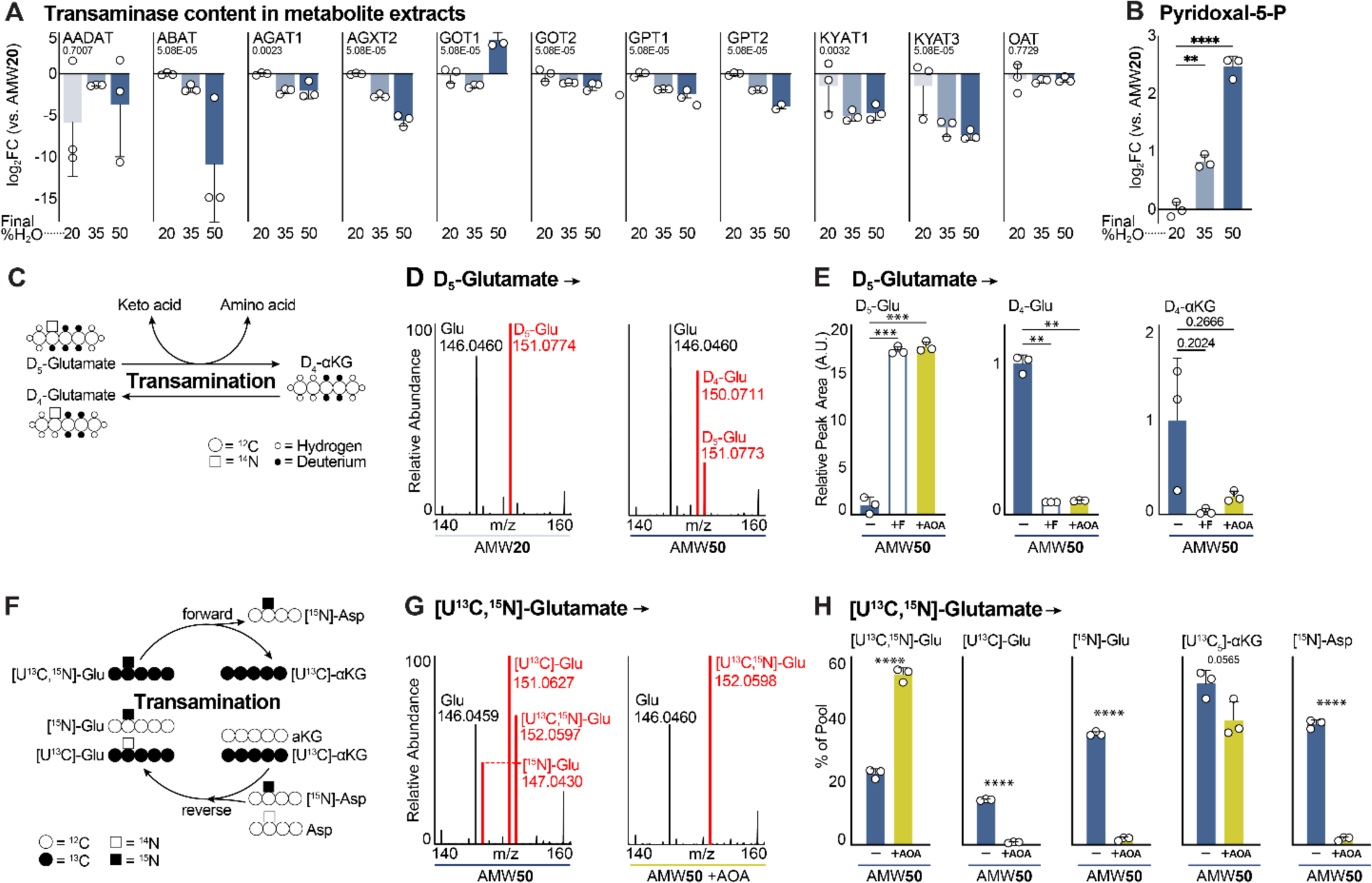
Active transaminase futile cycling in dried and resuspended metabolite extracts. (**A**) Relative abundance of murine liver transaminases in metabolite extracts from AMW20, AMW35, and AMW50 samples. Main effect FDR value shown (mean± SD, *n*=3 per group). (**B**) Relative abundance of pyridoxal-5-phosphate in murine liver AMW20, AMW35, and AMW50 samples. Significance calculated by Welch ANOVA (mean ± SD, *n*=3 per group). (**C**) Schematic showing deuterium labeling patterns for a transaminase reaction converting glutamate to α-ketoglutarate. (**D**) Mass spectra of glutamate in murine liver following D_5_-glutamate addition at resuspension. Representative AMW20 and AMW50 sample spectra are shown. Peaks corresponding to unlabeled glutamate (146.0460 m/z), D_4_-glutamate (150.0711), and D_5_-glutamate (151.0773 m/z) are shown. (**E**) Relative abundance of D_5_-glutamate-derived metabolites in murine liver. D_5_-glutamate was added at resuspension. Significance calculated by Welch’s ANOVA (mean ± SD, *n*=3 per group). (**F**) Schematic showing ^13^C and ^15^N labeling patterns for a cyclical transaminase reaction following [U^13^C,^15^N]-glutamate input. (**G**) Mass spectra of glutamate in murine liver following [U^13^C,^15^N]-glutamate addition at resuspension. Representative AMW50 and AMW50 +AOA sample spectra are shown. Peaks corresponding to unlabeled glutamate (146.0460 m/z), [^15^N]-glutamate (147.0430 m/z), [U^13^C]-glutamate (151.0627), and [U^13^C,^15^N]-glutamate (152.0598 m/z) and are shown. H) Relative abundance of [U^13^C,^15^N]-glutamate-derived metabolites in murine liver. [U^13^C,^15^N]-glutamate was added at resuspension. Significance calculated by Welch ANOVA (mean ± SD, *n*=3 per group). *p<0.05, **p<0.01, ***p<0.001, ****p<0.0001.

We next utilized doubly labeled [U-^13^C^15^N] glutamate to gain mechanistic insight into specific metabolic interconversions in AMW extracts. In this approach, amino acid formation from a transaminase reaction would only gain the ^15^N, and αKG would gain [U-^13^C] label. In the reverse reaction, [^15^N] labeled amino acids would donate the labeled nitrogen to unlabeled αKG to generate single labeled [^15^N] glutamate (**Figure 5F**). Note that reverse transaminase with [U-^13^C] αKG and a [^15^N] amino acid would lead to the regeneration of [U-^13^C^15^N] glutamate, which is indistinguishable from the originally added [U-^13^C^15^N] glutamate. Consequently, [^15^N] glutamate content underestimates Glu → αKG → Glu futile cycling. Separate [U-^13^C] and [^15^N] glutamate species were observed in AMW50 extracts and this was prevented by AOA addition (**Figure 5G**), positively demonstrating a transaminase futile cycle. AOA-mediated transaminase inhibition completely preserved initial [U-^13^C^15^N] glutamate and prevented the formation of [U-^13^C] glutamate and [^15^N] glutamate isotopologues (**Figure 5H**). [U-^13^C] αKG was not significantly different than control with AOA treatment. We interpret this as an indication of the presence of a non-AOA inhibitable deamination of [U-^13^C^15^N] glutamate through an undefined mechanism (**Figure 5I**). Finally, we evaluated ^15^N incorporation into the amino acids that were detected in our studies (**Figure S6**). Significant nitrogen labeling was found only in aspartate (**Figure 5I**), providing clear evidence of glutamate-aspartate transaminase activity in metabolite extracts. Collectively, these results provide clear evidence of enzymatic activity and metabolite interconversions in resuspended metabolomics extracts.

### A diverse proteome in metabolite extracts spans sample and extraction types

The complex interaction between extraction water content and metabolite abundance (**Figures 1-2**) appears to be driven, in part, by the hundreds of proteins contaminating AMW extracts (**Figure 4**) that can be enzymatically active (**Figure 5**). We next sought to understand whether this phenomenon was restricted to the sample type (liver) or extraction modality (AMW) that we selected. First, we evaluated the metabolomic and proteomic landscape of metabolite extracts from different sample types (mouse brain, skeletal muscle, perigonadal adipose tissue, plasma, and human HEK cells) extracted with AMW20, AMW35, and AMW50 metabolite extraction modalities. In all sample types, metabolite profiling revealed a prominent effect of extraction water content, with water-content separation by PCA occurring in either PC1(adipose, brain, skeletal muscle) or PC2 (HEK293, plasma) (**Figure 6A**, **Supplemental Table 3**). In all matrices, we observed an increase in ATP peak area with increased water content, though the magnitude of this increase was matrix dependent (**Figure S7A**). Plasma and skeletal muscle, but not other sample types, showed evidence of in-extract transaminase activity using D_4_/D_5_ glutamate as a readout (**Figure 7B**). Using proteomics, a diverse population of proteins were detected in all sample types and, like in the liver, this was strongly influenced by extraction water content (**Figure 6B** and **Supplementary Table 4**). Furthermore, GSEA of each type revealed metabolic gene sets within the top five enrichments for adipose tissue, HEK cells, and skeletal muscle (**Figure 6C**). Consistent with elevated D_4_/D_5_ Glu we also observed GOT1 and GOT2 in muscle (**Figure S7C**). Collectively, these data demonstrate pervasive protein contamination of metabolite extracts, and that the specific proteins enriched in the metabolite supernatant will vary between sample types.

**Fig. 6.**
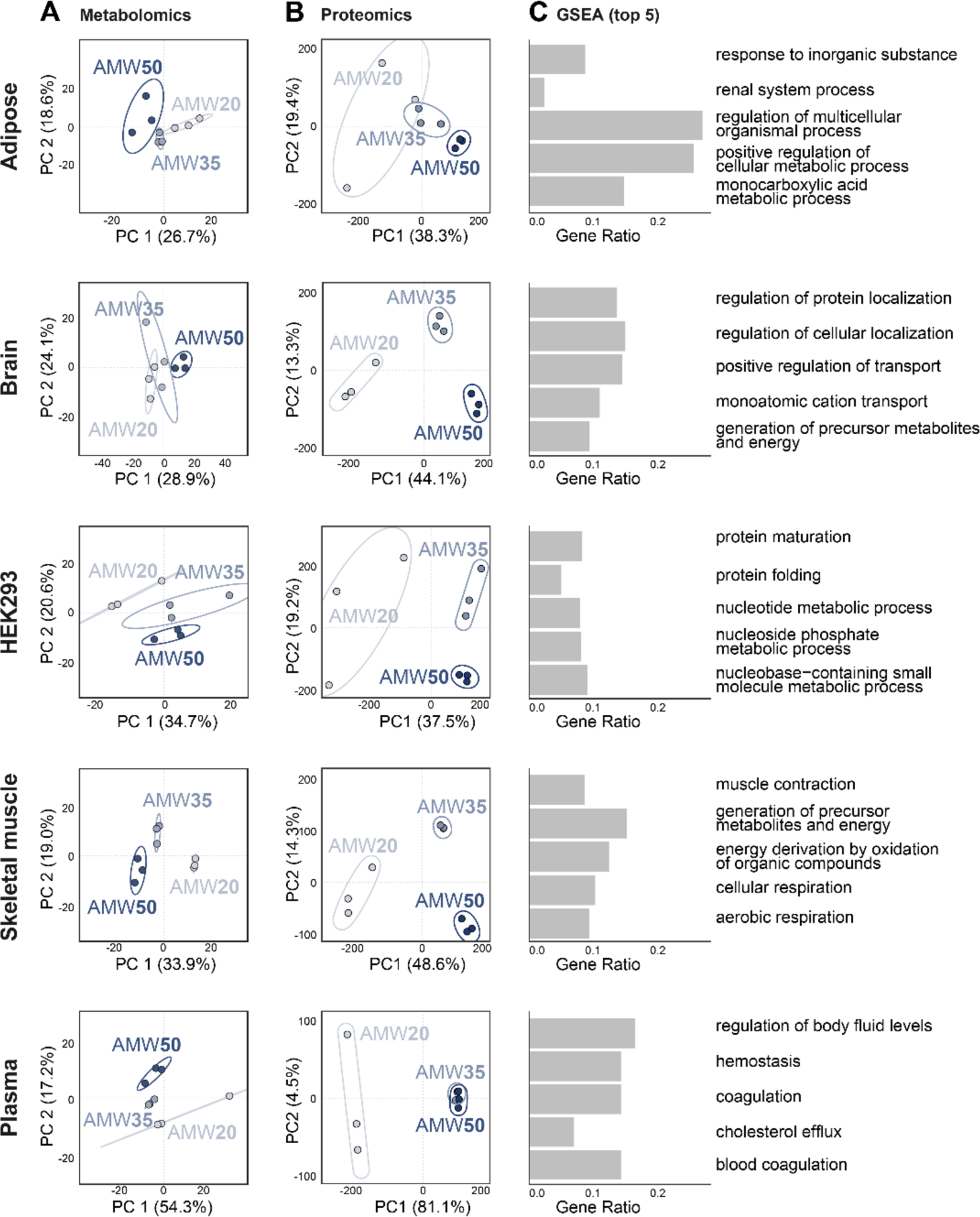
Metabolomic and proteomic content of metabolite extracts across tissue types. (**A**) PCA of AMW20, AMW35, and AMW50 metabolites across matrices. Matrices include murine liver, adipose, brain, muscle, and plasma, and human HEK293 cells (95% confidence ellipse, *n*=3 per group). (**B**) PCA of AMW20, AMW35, and AMW50 proteins across matrices. Matrices include murine liver, adipose, brain, muscle, and plasma, and human HEK293 cells (95% confidence ellipse, *n*=3 per group). (**C**) Top 5 most extraction water-content impacted biological processes based on FDR q-values within each matrix. Matrices include murine liver, adipose, brain, muscle, and plasma, and human HEK293 cells (*n*=3 per group).

**Fig. 7.**
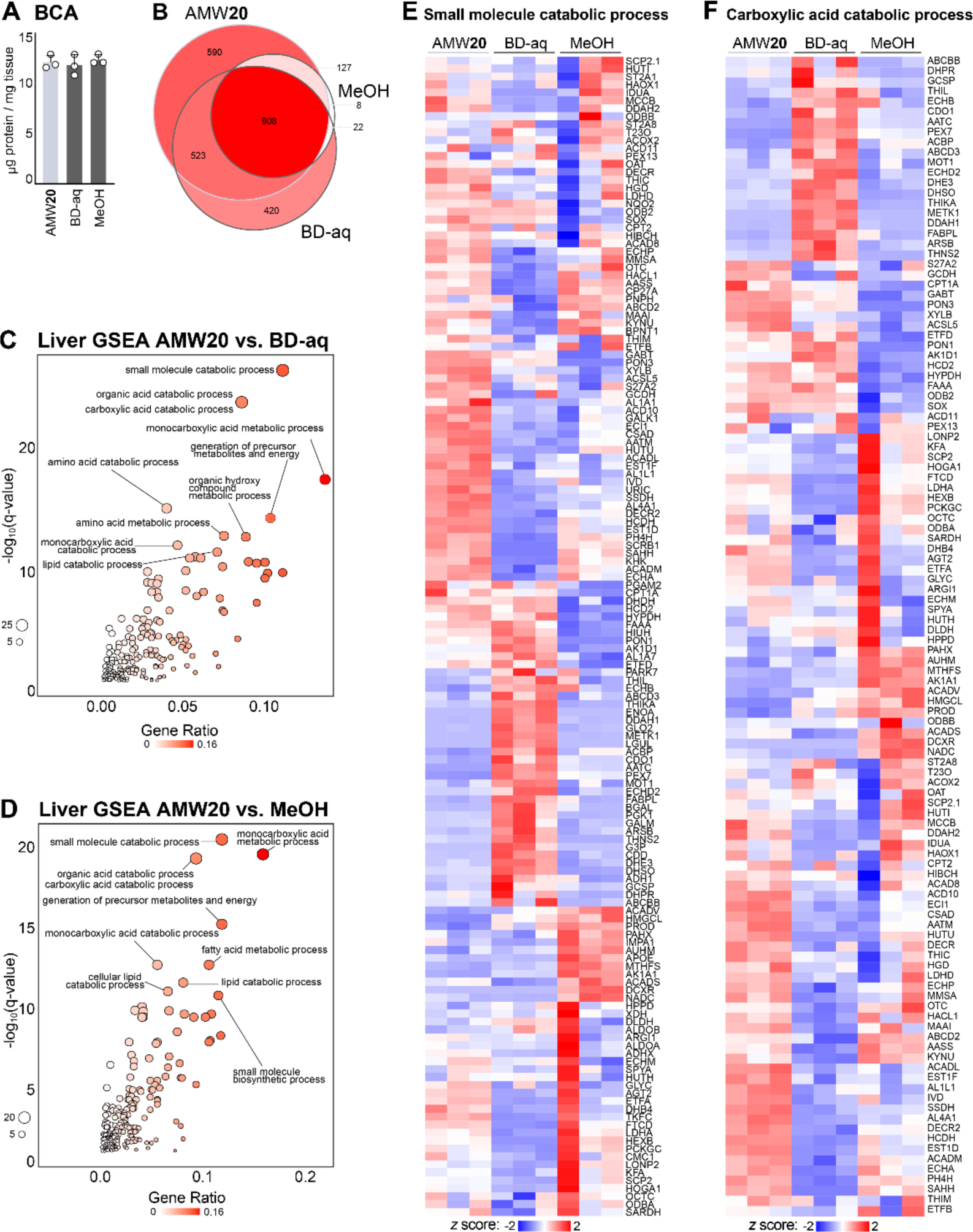
Proteomic composition of multiple metabolite extraction types. (**A**) BCA determination of protein content, normalized by starting tissue amount across AMW20, Bligh-Dyer aqueous phase (BD-aq), and 80% methanol extractions (*n*=3/group). (**B**) Number of protein groups identified in AMW20, Bligh-Dyer aqueous phase, and 80% methanol metabolite extractions (*n*=3/group). (**C**) GSEA BD-aq *versus* AMW20 Biological processes with the top 5 most impacted pathways based on FDR q-values, labelled. Points are sized by –Log_10_(q-value) to highlight significance and colored by the number of differentially abundant proteins in a given pathway divided by the number of proteins in that pathway (GeneRatio). (**D**) GSEA BD-aq *versus* AMW20 gene set enrichment analysis (biological processes) of proteins significantly impacted by extraction method. The top 5 pathways (based on FDR q-values) are labelled. Points are sized by –Log_10_(q-value) and colored by the ratio of differentially abundant proteins divided by total proteins in each pathway (GeneRatio). (**E**) GSEA MeOH *versus* AMW20 gene set enrichment analysis (biological processes) of proteins significantly impacted by extraction method. The top 5 pathways (based on FDR q-values) are labelled. Points are sized by –Log10(q-value) and colored by the ratio of differentially abundant proteins divided by total proteins in each pathway (GeneRatio). (**F**) Heatmap depicting relative abundance of small molecule catabolic process-associated proteins across extraction conditions and row standardized. All differentially abundant proteins (based FDR p-value) are shown. Significance calculated via LIMMA-eBayes on Log_2_ transformed proteins without any missing values. (**G**) Heatmap depicting relative abundance of carboxylic acid catabolic process-associated proteins across extraction conditions and row standardized. All differentially abundant proteins (based FDR p-value) are shown. Significance calculated via LIMMA-eBayes on Log_2_ transformed proteins without any missing values.

We next evaluated the proteomic composition of liver samples with two additional extraction types: 80% methanol (MeOH) and the metabolite containing aqueous phase of a Bligh-Dyer extraction (BD-aq; chloroform:methanol:water, 2:2:1.8 v/v) as in **Figure 4** and **6**. Protein content per unit mass of liver, as assessed by BCA, was consistent across extraction modalities (**Figure 7A**). Proteomics analysis revealed that a subset of 908 proteins was common to all three extraction types, with 590 being unique to AMW, 8 to MeOH, and 420 to BD-aq (**Figure 7B**). GSEA completed on proteins that were differentially abundant between extraction types revealed, as in (**Figure 4**), revealed the strong enrichment of metabolic pathways (**Figure 7C-D**). From GSEA alone, it was not clear whether these enrichments were being driven by metabolic proteins strongly abundant in one extraction type but lowly in others. To address this, we examined the relative abundance of proteins across extraction types in the two most enriched gene sets: “small molecule catabolic process” (**Figure 7E**) and “organic acid carbolic process” (**Figure 7F**). This revealed a complex interaction between extraction modality and metabolic protein abundance. Thus, metabolic protein carryover in metabolite extracts is not restricted to a specific extraction approach, but rather this is a ubiquitous phenomenon that is likely relevant to all metabolite extraction modalities.

### Untargeted metabolomics exposes other protein-mediated, post-extraction metabolic interconversions

Given the breadth of metabolic proteins contained in metabolite extracts (**Figures 4, 6** and **7**) and post-extraction transaminase activity (**Figure 5**), we sought to more comprehensively characterize metabolite interconversions caused by protein carryover. We hypothesized that post-resuspension enzymatic activity would manifest as time-dependent changes in metabolite levels. We further reasoned that complex proteome-metabolome interactions in extracts may yield metabolite products not annotated on our methods, prompting us to take an untargeted metabolomics approach. Unfiltered and filtered (F) AMW20 and AMW50 liver metabolite extracts were assessed through repeat injections over time (approximately every 2 hours for up to 84 hours) after resuspension by ESI negative-mode ion-paired LCMS. PCA of untargeted metabolomics data revealed distinct separation of the four groups (AMW20, AMW20F, AMW50, and AMW50F), but a time-dependent distribution in PC1 was prominently observed in unfiltered AMW50 extracts (**Figure 8A-B**).

**Fig. 8.**
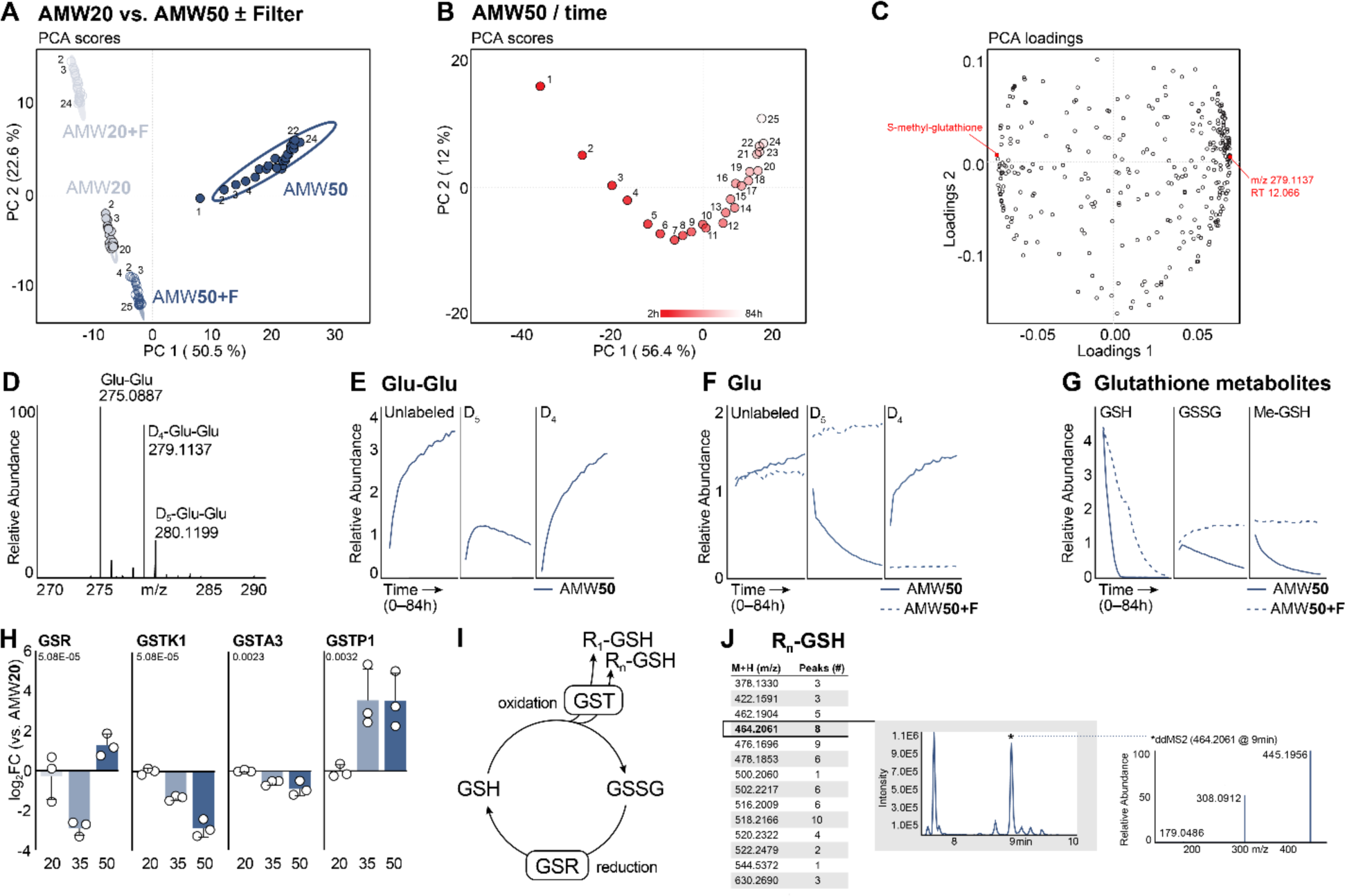
Untargeted metabolomics reveals time-dependent, protein mediated effects in dried and resuspended metabolomics extracts. (**A**) PCA of murine liver metabolites from AMW20 ±Filter and AMW50 ±Filter samples over time (25 repeat injections, 0-84 hr post-resuspension). (**B**) PCA scores of murine liver metabolites from AMW50 samples over time (25 repeat injections, 0-84 hr post-resuspension). (**C**) PCA loadings of murine liver metabolites from AMW50 samples over time (25 repeat injections 0-84 hr post-resuspension). Red dots represent the most positive and negative values along PC1 (+0.073, m/z 279.1137 at RT 12.066; no formula prediction and −0.073; S-methyl-glutathione m/z 320.0924 at RT 8.792; predicted chemical formula C_11_H_19_N_3_O_6_S Δppm = 1.0). (**D**) Mass spectra of glutamyl-glutamate (Glu-Glu) in murine liver following D_5_-glutamate addition at resuspension. Representative AMW50 sample spectrum is shown. Peaks corresponding to unlabeled Glu-Glu (275.0887 m/z), D_4_-Glu-Glu (279.1137 m/z), and D_5_-Glu-Glu (280.1199 m/z) are shown. (**E**) Relative abundance of D_5_-glutamate labeled glutamyl-glutamate (Glu-Glu) over time (25 repeat injections 0-84 hr post-resuspension). (**F**) Relative abundance of D_5_-glutamate-labeled glutamate over time (25 repeat injections 0-84 hr post-resuspension). (**G**) Relative abundance of GSH, GSSH, and Me-GSH over time (25 repeat injections 0-84 hr post-resuspension). (**H**) Relative abundance of murine GSR and GSTs in metabolite extract across AMW20, AMW35, and AMW50 extraction conditions. Main effect FDR value shown (mean +/− SD, *n*=3 per group). (**I**) Schematic showing GSR-mediated glutathione reduction, spontaneous glutathione oxidation, and GST-mediated glutathionylation. (**J**) Precursor ions with glutathione fragment ion, m/z = 308.0912 using the T3 method. Number of chromatographic peaks identified with a peak width greater than 6 s, and peak intensity 1E5 or greater. Representative chromatogram for precursor ion m/z464.2061, and the ddMS2 for that ion at RT 9.0 min.

Evaluation of PCA loadings (**Figure 8C**) for AMW50 showed this time-dependent effect was driven by many features, including an unknown compound (PC1 = +0.073, m/z 279.1137, RT 12.066 min; no formula prediction). Interestingly, this unknown compound coelutes with α-glutamyl-glutamate dipeptide (Glu-Glu; m/z 275.0886, RT 12.085 min). The mass difference between the unknown and Glu-Glu is 4.0256 Da, which is consistent with the mass shift expected by replacing four hydrogens (1.0078 Da) with deuterium (2.0141 Da) (**Figure 8D**). This suggests that the unknown compound is D_4_-labeled Glu-Glu, arising from the D_5_ Glu added at resuspension (**Figure 5**). Glu-Glu was completely absent in filtered extracts, whereas we noted a time dependent increase in unlabeled Glu-Glu in unfiltered extracts, indicating a that Glu-Glu is being generated de novo after resuspension (**Figure 8E)**. We also observed an initial increase in D_5_ Glu-Glu followed by a relative decrease as a fraction of the total Glu-Glu pool (**Figure 8E)**. Glu-Glu. This biphasic response of D_5_ Glu-Glu corresponds with the concomitant, time-dependent conversion of D_5_ Glu to D_4_ Glu (**Figure 8F**). Further, with D_4_ Glu and D_5_ Glu accounting for roughly 50% of the total free glutamate pool (**Figures 3, 5, 8F**), if Glu-Glu dipeptide production was sourced entirely from the free glutamate pool, then it should yield M+8 (D_4_Glu-D_4_Glu; [M-H] m/z = 283.1376), M+9 (D_4_Glu-D_5_Glu; [M-H] m/z = 284.1439), and M+10 (D_5_Glu-D_5_Glu; [M-H] m/z = 285.1501) deuterium isotopologues that sum to roughly 25% of the total Glu-Glu pool. Instead, none of these higher order deuterium (M+8, M+9, M+10) isotopologues were observed (**Figure 8D**). This suggests that only one glutamate in the Glu-Glu dipeptide arises from the free glutamate pool while the other arises from another glutamate source, such as protein hydrolysis, and that it is likely enzymatic since Glu-Glu formation is completed prevented by protein removal.

Another compound noted from our untargeted metabolomics study was putatively identified as S-methyl-glutathione (**Figure 8C**; PC1 = −0.073; m/z 320.0924, RT 8.792 min; predicted chemical formula C_11_H_19_N_3_O_6_S, Δppm = 1.0), which decreased over time (**Figure 8G**). This led us to examine reduced (GSH) and oxidized (GSSG) glutathione pools. In AMW50 and AMW50F extracts (**Figure 8G**), we observed a time-dependent decrease in GSH, likely due to spontaneous oxidation, and a concomitant rise in GSSG. Non-enzymatic degradation of GSH through cysteinyl-glycine bond cleavage can be ruled out as previous NMR-based studies demonstrate this reaction exhibits slow kinetics and occurs on the order of weeks [53]. Interestingly, GSH depletion occurred more quickly in AMW50 extracted samples but was not met with a corresponding rise in GSSG (**Figure 8G**). Instead, GSSG, like Me-GSH, also decreased over time. The decrease in GSSG in an oxidizing environment suggests an enzymatic mechanism, and indeed our proteomics data revealed the presence of twenty-five glutathione metabolizing enzymes (**Figure S8**). Included in these is glutathione reductase (GSR) and sixteen glutathione S transferases (**Figure 8H**, **Figure S8**). Thus, the potential exists for GSSG reduction back to GSH and subsequent electrophilic conjugation to a variety of compounds and/or proteins (**Figure 8I)**.

Along these lines, we examined untargeted ddMS2 spectra from reversed phase (T3) LCMS in AMW50 and AMW50F samples for precursor ions with the characteristic glutathione fragment ion (m/z = 308.0912). We identified 14 precursor ions that gave rise to 67 distinct isobaric (+/− 2 ppm) chromatographic peaks (peak width greater than 6sec, and peak intensity 1E^5^ or greater) in AMW50 samples that were absent in AMW50F samples (**Figure 8J**). Interestingly, we noted several precursor ions that differed in m/z by 2.0157Da. This is the same mass difference that occurs as acyl chains gain double bonds, losing two hydrogens (e.g., 18:0 fatty acid [M+H] = 285.2788 m/z and 18:1 fatty acid [M+H] = 283.2632 m/z, Δm/z = 2.0156). This observation, considered together with the strong retention observed in reversed phase chromatography and the large number of isobaric peaks, supports the hypothesis that lipids are being gluthathionylated in metabolite extracts. Though the molecular identities of these glutathionylated compounds remain unknown, these data suggest the presence of a protein-mediated glutathione sink that explains the time-dependent, in-extract loss of total glutathione (**Figure 8G**) in AMW50 samples.

### Protein removal by filtration in combination with high-water extraction content improves metabolomics coverage

Increasing extraction water content improves the recovery of hydrophilic compounds, such as nucleotides (**Figure 1**). However, this exposes samples to post-extraction enzymatic activity and associated confounding effects that can be prevented by removing proteins with 3 kDa filtration (**Figures 3-5, 8**). So, we hypothesized that combining high-water AMW metabolite extraction with 3kDa filtration would simultaneously improve compound recovery while mitigating the risks associated with protein carryover. We subjected liver samples to AMW20 and AMW50 extractions with or without 3 kDa filtration. Targeted metabolomics was performed on three orthogonal methods as described in **Methods**. Of the 287 compounds detected, 212 were significantly different by ANOVA and *post hoc* multiple comparisons test. Consistent with the water titration experiment (**Figure 1B, D**). Hierarchical clustering revealed distinct clades of compounds that were affected by filtration (**Figure 9A and Figure S9A**). This included elevated levels of polar metabolites, such as nucleotides, in AMW50 that were unaffected by filtration (**Figure S9B**). Interestingly, filtration depleted long-chain lipid species including acyl-carnitines (**Figure 9A, Cluster 1, and Figure S9C**) in the AMW50 samples, revealing an extraction water by filter interaction. This is likely a non-specific binding of lipids to the filter material (regenerated cellulose membranes), rather than a size exclusion. Another cluster containing many amino acids was elevated in AMW50F *versus* AMW50 (**Figure 9A**, Cluster 2), suggesting that these compounds are lost through a protein-mediated mechanism in AMW50 extracts. A third cluster revealed compounds that are elevated in AMW50, but prevented by filtration (**Figure 9A**, Cluster 3), including glutamyl-glutamate (**Figure 8E**). Interestingly, several nucleosides and nucleobases including adenine, uracil, and adenosine are constituents of Cluster 2 (**Figure 9B**) while hypoxanthine, inosine, guanosine, uridine, and xanthosine are grouped in Cluster 3 (**Figure 9C**). This bidirectional response of metabolic neighbors in AMW50 that is prevented by filtration may suggest pyrimidine/purine metabolic enzymes as another set of in-extract active proteins. Similarly, pyridoxal content was diminished (**Figure 9B**) while pyridoxal 5P was elevated (**Figure 9C**) in AMW50 extracts, which was prevented by filtration. This apparent protein-mediated, in-extract conversion of pyridoxal to pyridoxal-5P may contribute to transaminase activity in these extracts (**Figure 5**). Collectively, these data support the use of high-water AMW metabolite extraction with 3 kDa filtration to improve polar metabolite recovery while preventing enzyme mediated metabolic interconversions. However, we note that this approach would require further optimization to be suitable for analyzing polar lipids.

**Fig. 9.**
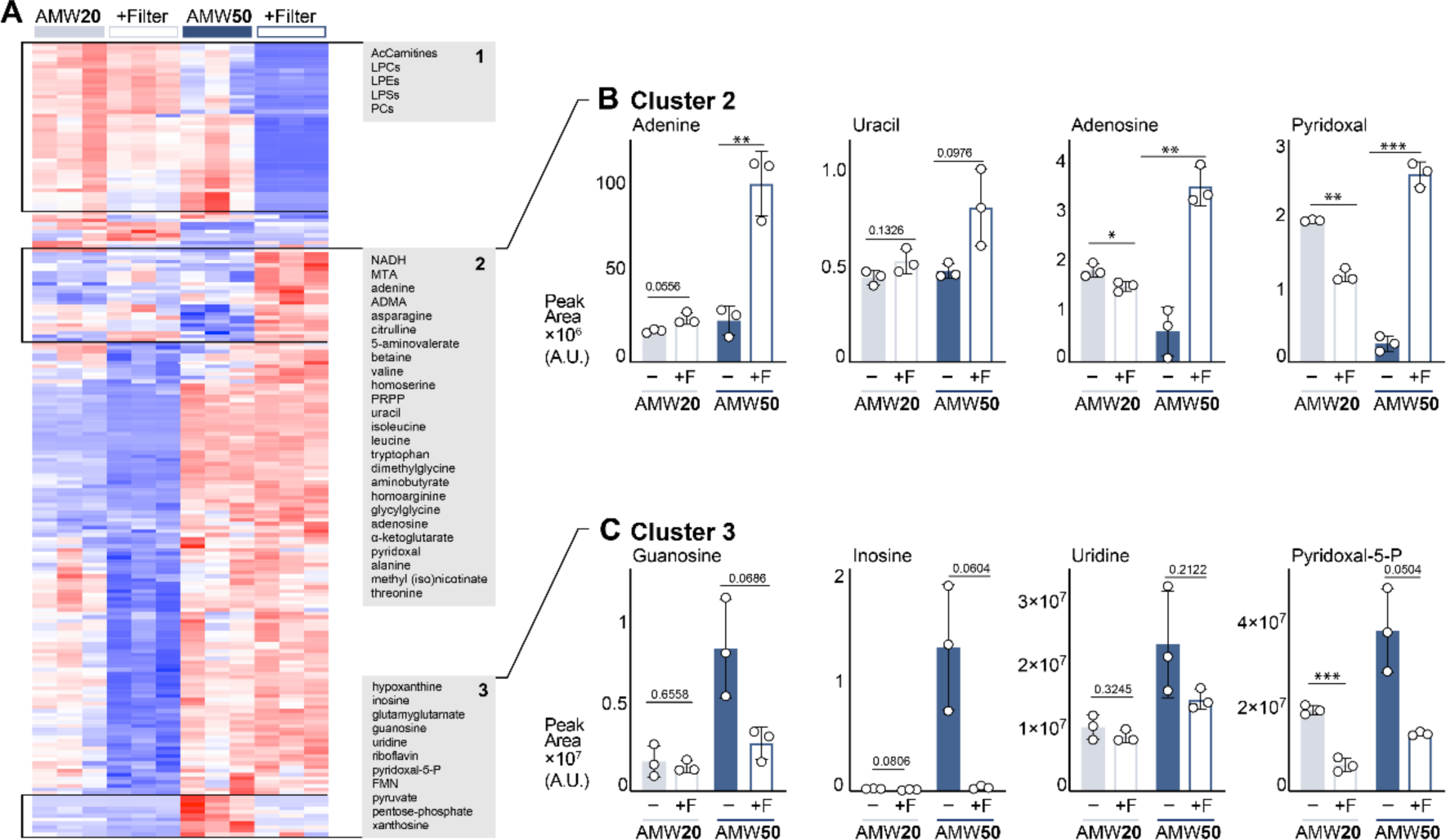
3kDa filtration in high water content AMW extracts improves metabolite recovery and prevents unwanted protein-mediated effects. (**A**) Heatmap depicting relative abundance of murine liver metabolites across AMW20 ± Filter and AMW50 ± Filter extraction conditions. All significantly different metabolites are shown. Significance was calculated by ANOVA using MetaboAnalyst 5.0, and row hierarchical clustering was performed using Morpheus (Broad). (**B**) Relative abundance of Cluster 2-related murine liver metabolites across AMW20 ± Filter and AMW50 ± Filter extraction conditions. Significance calculated by Welch’s ANOVA (mean ± SD, *n*=3 per group). (**C**) Relative abundance of Cluster 3-related murine liver metabolites across AMW20 ± Filter and AMW50 ± Filter extraction conditions. Significance calculated by Welch ANOVA (mean ± SD, *n*=3 per group).

## Discussion

Here, we present the first thorough characterization of the protein composition of metabolomics extracts across common extraction methods and sample types. We demonstrate that metabolite extract proteomes are enriched for metabolic enzymes that retain their activity despite exposure to denaturing chemical conditions. Our findings have important implications, as post-extraction enzymatic activity poses a threat to accurate biological phenotype detection and characterization. Finally, we address this previously unquantified, in-extract proteome by presenting a novel extraction method that expands compound coverage and incorporates passive 3 kDa filtration to remove proteins.

Metabolite extraction is the critical first step in metabolomics workflows as it defines the analytical scope of downstream chromatography-mass spectrometry analysis. The use of 80% polar solvents is widely used for profiling metabolomics studies. This originated with an 80% methanol (aqueous) with EGTA extraction that preserved and maximized recovery of nucleotides [26]. This polar extraction method was refined to 40% methanol, 40% acetonitrile, and 20% water with formic acid for enhanced *E. coli* metabolite extraction [27], but formic acid was later determined to be dispensable [23]. This AMW method is widely used for metabolomics studies (see [38–50] for a non-exhaustive list) and was the basis of the current approach. However, despite its widespread use, AMW extraction water content varies widely in the literature [16, 22, 25, 54–58] and has not been systematically evaluated on a whole-metabolome scale. With this as motivation, we titrated extraction water content and observed a strong and complex metabolomic response. As an example, increasing extraction water content from 20% to 25% induced a 4-fold increase in detected nucleotide triphosphate peak area; this nucleotide effect plateaued at an ∼8-fold increase by 40% water. These data underscore the need to tightly control extraction water content, including the solvent:sample ratio, by adjusting extraction solvent volume to sample amount (i.e., tissue mass, cell number, volume, etc.) to account for sample water content. Indeed, we discovered that many metabolites responded to increased extraction water content in a manner not consistent with their intrinsic hydrophobicity, leading us to consider a protein-mediated effect occurring post-extraction.

The dogmatic view of metabolite extraction with polar solvents is that larger biomolecules such as RNA, DNA, and protein are insoluble. Indeed, the insoluble fraction can be subsequently extracted and analyzed, enabling single-sample multi-‘omic workflows [29–36]. However, the presence of these molecules in the insoluble fraction does not necessitate their absence from the soluble, metabolite containing fraction. The present work demonstrates that 2-4% of the total sample protein content remains and comprises over 1,000 individual proteins that are enriched for metabolic annotations. Moreover, these proteins are present in common metabolite extraction modalities including AMW20, 80% methanol, and Bligh-Dyer, though specific protein composition varies in each. Similarly, broad protein carryover was observed across sample types following AMW20 extraction. Collectively, our data demonstrate that metabolites (i.e., potential substrates) and proteins are being co-incubated during extraction and post-resolubilization.

Protein carryover in metabolite extracts can lead to post-extraction enzymatic activity and metabolite interconversions (**Figures 3** and **8**). Additionally, a diverse protein-metabolite interactome has been described [59] and it is unknown to what extent *de novo* metabolite-protein interactions will form. Metabolites that become bound to residual proteins will be masked from LCMS detection. Thus, both post-extraction enzymatic activity and metabolite-protein interactions pose a threat to faithful detection of biological phenotypes. Since the proteome and metabolome are unique to every extraction modality and sample type (**Figures 6** and **7**), the potential for confounding influence will be different in every scenario and therefore difficult to test, isolate, and observe. Thus, we posit that the prudent course is to take additional steps to ensure the removal of proteins from metabolite extracts.

Here, we elected to use 3 kDa filtration to remove proteins from metabolite extracts while preserving the metabolome. Our proteomics and metabolomics data supports the effectiveness of this approach. A similar approach using a 10 kDa filter was used on dried and resuspended extracts for NMR analysis, citing metabolite instability, with mixed results [60, 61]. This inconsistency may be explained considering our present result that enzymatic activity persists in resuspended extracts. Instead, we subjected the soluble fraction of the crude extract to 3 kDa filtration, then dried the eluate for metabolomics. 3 kDa filtration enabled higher aqueous solvent composition and ultimately improved metabolomic coverage while preventing in-extract metabolic conversions. However, this approach adds an extra step to the workflow that takes additional time, can introduce another source for variation, and potential sample loss. Despite these disadvantages, the 3 kDa filtration is the least obtrusive and best preserves the metabolome when compared to other protein removal options. Sample heating is a common approach to denature proteins but will have broad effects on metabolites. For example, glutamine cyclizes to pyroglutamate when moderately heated [62]. Sample acidification is another potential option, but one that will also affect the metabolome, and like polar-solvent precipitation, the completeness of protein precipitation is assumed rather than based on empirical evidence.

Another important implication of this work is in the potential for post-extraction changes in the isotopologue distribution from stable isotope tracing studies, obscuring biological interpretation. This is apparent in our observations of a D_5_-D_4_ glutamate transition that occurred in the absence of changes to the overall glutamate pool (**Figures 3, 5**). In this case, in-extract glutamate transamination leads to aspartate and α-ketoglutarate formation (**Figure 5**). This, in turn, mixes carbons between Glu and αKG, and nitrogen between Glu and Asp. If occurring in biological tracing experiments, such in-extract transamination could lead to false interpretations. This risk extends beyond the glutamate transamination observed here, as any post-extraction enzymatic activity may lead to tracer mixing between metabolic neighbors. Such activity may not be apparent in metabolite abundance alone and will likely differ across experimental approaches. Thus, 3kDa extract filtration is an attractive option to ensure protein removal and preservation of in vivo labeling patterns from stable isotope labeling studies.

Metabolite extract supernatants are commonly collected and dried in a vacuum evaporator, under a stream of nitrogen gas, or through lyophilization, then resuspended for analysis. This serves to both concentrate the extracts and to resuspend them in solvent compatible with the analytical approach. For LCMS workflows, resuspension solvent type should match the beginning mobile-phase composition of the chromatography. Thus, for reversed-phase applications where the chromatography gradient ramps from high-aqueous to high-organic composition, samples are commonly resuspended in water. In contrast, metabolomics-variations of normal phase methods, including hydrophilic-interaction (HILIC) and amide chromatography methods, where the gradient is organic-to-aqueous, samples are resuspended in a mixture of water and solvent (e.g., 50:50 acetonitrile:water) to make the sample compatible with the chromatography while concomitantly resolubilizing high-polarity metabolites of interest. In the present study, our observations of enzymatic activity in resuspended extracts were made exclusively in samples resuspended in water for reversed phased analysis. It is unclear whether enzymatic activity occurs using other resuspension solvents.

The presence of intact proteins in metabolomics samples likely has other practical consequences. Metabolomics methods rely heavily on retention time for compound identification, which makes chromatography stability and reproducibility foundational. The unintended introduction of protein to the analytical system may negatively affect the retention mechanisms of the column, leading to RT shifts, and/or protein buildup at the column inlet may cause pressure increases, peak broadening, and shortening column life. If these proteins elute from the column, then they may interfere with analyte detection through ion suppression or adduct formation. These issues are difficult to evaluate systemically, but removal of unwanted contaminates from analytical chemistry workflows is a guiding principle and one that is relatively easily accomplished in this case through 3 kDa filtration.

Our data offers mechanistic insight into the mathematical correction of time-dependent, analyte specific signal drift using periodic injections of pooled samples, which is common practice in metabolomics data processing [63–66]. Given instrument signal stability, changes over time in signal responses of individual metabolites are postulated to be compound intrinsic or otherwise enigmatic. For example, loss of a metabolite over time is thought to be due to compound instability, whereas signal gain over time may be attributed to longer solubilization in the resuspension solvent. Our data provide another possible mechanism for this phenomenon, that enzymatic activity and/or protein-metabolite binding kinetics may account for time-dependent compound signal drift. In such cases, the removal of proteins by 3 kDa filtration of extracts would stabilize the signal and obviate the need for peak area correction of these metabolites.

Here, we have discovered a diverse proteome in metabolite extracts, the scope of which is determined by a complex interaction between extraction modality and sample type. We further show that in-extract proteins can be active and drive post-extraction metabolome changes. This constitutes a previously unknown observer effect in metabolomics, putting biological phenotype detection at risk. We provide a practical solution to mitigate this problem by using 3 kDa filtration. Our novel workflow integrates increased extraction water content with filtration for protein removal and enhanced broad-coverage metabolomics.

## Materials and Methods

### Tissue collection and processing

Wild-type mice (C57B6J) of mixed ages and both sexes were used in this study. Mice were anesthetized with an isoflurane vaporizer and whole blood was collected from cardiac puncture into a 1.5 mL tube containing 10 µL of 0.5 M EDTA. Liver, epigonadal adipose tissue, gastrocnemius skeletal muscle, and brain, were then collected. Tissues were immediately snap frozen in liquid nitrogen (LN_2_) and stored at –80°C for later processing. Frozen tissues were pooled by type and pulverized with a mortar and pestle in LN_2_. Cryopulverized pooled tissues were thoroughly mixed, and, careful to avoid sample thawing, weighed into 30-40 mg aliquots into LN_2_ pre-chilled 2 mL bead mill homogenizer tubes (19-627, Omni). Whole blood with EDTA was kept on wet ice until processing (<60min) and centrifuged at 4,000 × *g* for 10 min at 4 °C. Plasma from each mouse was pooled, mixed by vortex, split into 40 µL aliquots, and stored at –80 °C. All mouse procedures were approved by the Van Andel Institute Institutional Animal Care and Use Committee.

### Cell culture

Human kidney epithelial HEK293 Phoenix-AMPHO cells (ATCC CRL-3213), were maintained in Dulbecco’s modified eagle medium (DMEM) no phenol red (Gibco), 10% fetal bovine serum (FBS), 5 mL penicillin-streptomycin (Gibco). Cells were maintained at 37°C in a 5% CO_2_ incubator. For metabolite extraction, cells were seeded at a density of 1×10^6^ per well onto 6-well plate wells. Cells were grown in DMEM, no phenol red (Gibco), 10% FBS, and 5 mL penicillin-streptomycin (Gibco) for 48 hr at 37°C in a 5% CO_2_ incubator. Cells were then washed twice with ice-cold NaCl, and plates were frozen on dry ice, and transferred into a –80°C freezer to be stored until extraction.

### Metabolite extractions

The efficiency of the tissue homogenization is affected by the fractional volume of the sample in the bead mill homogenization tube. Thus, all samples were homogenized using a ratio of 40 mg tissue/mL of solvent to ensure the same degree of homogenization among samples. When testing higher water concentrations, additional water was added after initial AMW20 (40% acetonitrile:40% methanol:20% water) homogenization to achieve the desired final water content (**Supplemental Table S1**). For consistency, this same approach was followed for sample types (cells, plasma) not requiring bead mill homogenization. Homogenization for Bligh-Dyer (2:2:1.8 chloroform:methanol:water) and 80% MeOH extractions were similarly performed at 40 mg/mL. In all extraction types, the same amount of tissue/cell/plasma equivalents was dried and processed for bicinchoninic acid assay (BCA) protein content, proteomics, or metabolomics as indicated.

Metabolites were extracted using one of the following approaches as indicated in text and figure legends. For AMW extraction, ice-cold acetonitrile (ACN, A955-4, Fisher), methanol (MeOH, A456-4, Fisher), and H_2_O (W6-4, Fisher) were added to each matrix at a 4:4:2 (v/v/v) ratio. For 80% MeOH extraction the approach was the same as AMW. For Bligh-Dyer extracts, samples were homogenized in ice-cold 1:1 chloroform (1024441000, Millipore Sigma):MeOH, followed by the addition of 0.9 volumes of water to achieve the final 2:2:1.8 ratio as reported previously [10].

For solid, tissue matrices, extraction solvent volume was determined by (sample mg)/40 mg x (mL solvent). For the Phoenix-AMPHO cell extraction, solvent volume was determined by (1000 μL /smallest sample cell number) x (sample cell number). For plasma extraction, solvent volume was determined by (800 μL/40 μL) x (sample μL). After solvent addition, extracts were either homogenized for 30 s (tissue) or vortexed for 10 s (biofluids, cells), sonicated for 5 min, and incubated on wet ice for 1 hr. For AMW extracts with greater than 20% water, after the initial 1 hr incubation additional water was added a final water percentage between 25-60% as indicated. These AMW+H_2_O samples were vortexed, sonicated for 5 min, incubated on wet ice for an additional 10 min. Following incubation, AMW20 and AMW +H_2_O samples were centrifuged at 17,000 × *g* and 4°C for 10 min. The supernatants were collected and centrifuged a second time at 17,000 × *g* and 4°C for 10 min to ensure complete precipitate clearance from the supernatant. 16 mg-tissue-equivalents of aqueous phase supernatant was collected and dried in a vacuum evaporator. In some cases, 8 mg equivalents were dried instead, but resuspension volume was also halved resulting in equivalent tissue concentration in resuspended extracts and on column. The amount of sample equivalents on column was tightly controlled throughout the study.

### pH analysis metabolite extracts

Dried murine liver metabolite extracts from AMW20-AMW60 extraction conditions were resuspended in H_2_O, vortexed, sonicated for 5 min, and incubated on wet ice for 10 min. Resuspended sample extracts (50 μL) were transferred to a Seahorse XF-96 assay plate (101085-004, Agilent) along with pH standards: pH 2 (SB101 + NaOH, Fisher Chemical), pH 4 (SB101, Fisher Chemical), pH 7 (SB107, Fisher Chemical), and pH 11 (SB115, Fisher Chemical). Samples and standards pH values were measured using a Seahorse XF-96, which is an accurate micro-pH meter down to 25 μL (Figure S4). The amount of sample equivalents per resuspension extract was tightly controlled across samples.

### Protein removal from metabolite extracts by 3 kDa filtration

Amicon Ultra-2 3K centrifugal filter devices (UFC200324, Millipore) were pre-rinsed with 1.0 mL of LCMS grade water, and centrifugated for 20 min at 4,000 × *g*. All eluate and remaining unfiltered water were removed and immediately followed by the addition of metabolite supernatant to prevent membrane drying. Following the initial centrifugation of metabolite extraction (above), supernatant was transferred to the centrifugal filter device, and centrifuged at 4°C for 60 min at 4,000 × *g*; 8 mg-tissue-equivalents (determined by % volume of initial extraction volume) of eluate was collected and dried in a vacuum evaporator.

### Protein extraction from dried metabolomics samples

Dried metabolomic supernatant extracts were processed using the EasyPep Mini MS Sample Prep Kit (A40006, Thermo Fisher Scientific). Briefly, dried metabolite extract supernatants were resuspended in 100 µL of lysis solution per 16 mg tissue equivalents to extract proteins. Proteins were quantified using the Pierce BCA Protein Assay Kit (23227, Thermo Fisher Scientific). Proteins were reduced and alkylated at 95°C for 10 min, and samples were digested overnight with Trypsin/Lys-C at 30°C at a ratio of 10:1 (protein:enzyme (w/w)). Samples were cleaned with kit supplied peptide clean up columns and dried down in a Genevac SpeedVac prior to resuspension for instrument analysis. Samples were resuspended in 50 µL 0.1% formic acid (FA) (LS118-1, Fisher Scientific) and diluted with 50 µL of 0.1% trifluoroacetic acid (TFA) (LS119-500, Fisher Scientific).

### Preparation of liver samples for proteomics

Tissue samples were homogenized on the Bead Ruptor Elite (19-042E, Omni International) for 30 s in protein lysis solution. Samples were sonicated and clarified via centrifugation and transferred to a Protein LoBind Eppendorf tube. Proteins were quantified using the Pierce BCA Protein Assay Kit (23227, Thermo Fisher Scientific).100 µg of protein was aliquoted to a 1.5 mL screw top tube (72.703.600, Sarstedt) and digested using the EasyPep Mini MS Sample Prep Kit (A40006, Thermo Fisher Scientific). Briefly, sample volume was adjusted to 100 µL with lysis solution, proteins were reduced and alkylated at 95°C for 10 min, and samples were digested overnight with Trypsin/Lys-C at 30°C at a ratio of 10:1 (protein:enzyme (w/w)). Samples were cleaned up with kit supplied peptide clean up columns and dried down in Genevac SpeedVac prior to resuspension for LC-MS/MS. Samples were resuspended in 50 µL 0.1% FA (LS118-1, Fisher Scientific) and diluted with 50 µL of 0.1% TFA (LS119-500, Fisher Scientific).

### Protein Quantitation

Proteins were quantified using the Pierce BCA Protein Assay Kit (23227, Thermo Fisher Scientific) following the vendor supplied protocol. Samples were diluted in LCMS H_2_O (W6-4, Fisher Scientific). Plates were read at an absorbance of 562 nm using the Synergy LX Multi-Mode Reader and Gen5 software was used for data analysis (BioTek/Agilent). Polynomial regression was used in the Gen5 software to calculate protein concentrations to a protein standard curve after an average blank absorbance subtraction.

### Data-independent Acquisition (DIA) LC-MS/MS Proteomics

DIA analyses were performed on Orbitrap Eclipse coupled to Vanquish Neo system (Thermo Fisher Scientific). The FAIMS Pro source (Thermo Fisher Scientific) was located between the nanoESI source and the mass spectrometer. 2 μg of digested peptides were separated on a nano capillary column (20 cm × 75 μm I.D., 365 μm O.D., 1.7 μm C18, CoAnn Technologies, Washington, # HEB07502001718IWF) at 300 nL/min. Mobile phase A consisted of LC/MS grade H_2_O (LS118-500, Fisher Scientific), mobile phase B consisted of 20% LC/MS grade and H_2_O and 80% LC/MS grade acetonitrile (LS122500, Fisher Scientific), and both mobile phases contained 0.1% FA. The LC gradient was: 1% B to 26% B in 51 min, 80% B in 5 min, and 98% B for 4 min, with a total gradient length of 60 min. For FAIMS, the selected compensation voltage (CV) was applied (−45V and −65V) throughout the LC-MS/MS runs. Full MS spectra were collected at 120,000 resolution (full width half-maximum; FWHM), and MS2 spectra at 30,000 resolution (FMWH). Both the standard automatic gain control (AGC) target and the automatic maximum injection time were selected. A precursor range of 380-980 m/z was set for MS2 scans, and an isolation window of 50 m/z was chosen with a 1 m/z overlap for each scan cycle. 32% HCD collision energy was used for MS2 fragmentation.

#### DIA Database Search

DIA data was processed in Spectronaut (version 18, Biognosys, Switzerland) using direct DIA. Data was searched against *Mus musculus* or *Homo sapiens* reference proteomes as appropriate. The manufacturer’s default parameters were used. Briefly, trypsin/P was set as digestion enzyme and two missed cleavages were allowed. Cysteine carbamidomethylation was set as fixed modification, and methionine oxidation and protein N-terminus acetylation as variable modifications. Identification was performed using a 1% q-value cutoff on precursor and protein levels. Both peptide precursors and protein false discovery rate (FDR) were controlled at 1%. Ion chromatograms of fragment ions were used for quantification. For each targeted ion, the area under the curve between the XIC peak boundaries was calculated.

### LC/MS Metabolomics

#### Metabolomics approach

Dried metabolomics extracts were first resuspended in 100% water containing 25 µg/mL D_8_-Tryptophan (DLM-6903-0.25, Cambridge) for analysis using an ion-paired reversed phase chromatography on an Orbitrap Exploris 240 in ESI negative mode (described below). When appropriate for experimental design, final concentrations of 25 µg/mL D_5_-Glutamate (DLM-556, Cambridge), 25 µg/mL U-C^13^N^15^ Glutamate (CNLM-554-H-0.25, Cambridge), and 0.5 mM aminooxyacetic acid (28298, Cayman) were also added to the resuspension solvent. Resuspension volumes were varied by sample type to achieve uniform sample-equivalents per volume. This was 80 µg/µL for tissue, 160 µL/µL for plasma, and 6.70E+3 cells/µL. 2 µL of resuspended sample were injected on column for each method below. For deeper metabolomic coverage, samples were again dried and resuspended in 50% ACN (v/v) and analyzed using two orthogonal chromatographies (Waters BEH Amide and T3, described below) on an Orbitrap ID-X in ESI positive mode. For all experiments, process blanks were analyzed before and after experimental samples. Pooled samples were injected twice before experimental samples for column conditioning, and every 6-10 sample injections thereafter. Data dependent MS2 (ddMS2) data was collected from pooled samples for compound ID. In some experiments, ddMS2 was also collected on experimental replicates to identify group-specific compounds.

#### Tributylamine ion paired reversed phase LC/MS

As previously reported [9, 10, 36, 67], mobile phase A was LC/MS H_2_O (W6-4, Fisher) with 3% LC/MS grade MeOH (A456-4, Fisher), mobile phase B was LC/MS grade methanol (A456-4, Fisher), and both mobile phases contained 10 mM tributylamine (90780-100ML, Sigma), 15 mM acetic acid, and 0.01% medronic acid (v/v, 5191-4506, Agilent). For the re-equilibration gradient, mobile phase A was kept the same, and mobile phase B was 99% LC/MS grade acetonitrile (A955-4, Fisher). Column temperature was kept at 35°C, flow rate 0.25 mL/min, and the solvent gradient was as follows: 0-2.5 min held at 0% B, 2.5-7.5 min from 0% B to 20% B, 7.5-13 min from 20% B to 45% B, 13-20 min from 45% B to 99% B, and 20-24 min held at 99% B. The analytical solvent gradient was followed by a 16 min re-equilibration gradient to prep the column before the next sample injection that went as follows: 0-0.05 min held at 99% B at 0.25 mL/min, 0.05-1 min from 99% B to 50% B and 0.25 mL/min to 0.1 mL/min, 1-11 min held at 50% B and 0.1 mL/min, 11-11.05 min from 50% B to 0% B at 0.1 mL/min, 11.05-14 min held at 0% B at 0.1 mL/min, 14-14.05 min held at 0% B and increased flow rate from 0.1 mL/min to 0.25 mL/min, and 14.05-16 min held at 0% B and 0.25 mL/min. Data were collected on an Orbitrap Exploris 240 using a heated electrospray ionization (H-ESI) source in ESI negative mode. The mass spectrometer acquisition settings were as follows: source voltage - 2,500V, sheath gas 60, aux gas 19, sweep gas 1, ion transfer tube temperature 320°C, and vaporizer temperature 250°C. Full scan data were collected with a scan range of 70-800 m/z at a mass resolution of 240,000. Fragmentation data was collected using a data-dependent MS2 (ddMS2) acquisition method with MS1 mass resolution at 120,000, MS2 mass resolution at 15,000, and HCD collision energy fixed at 30%.

#### Amide and T3 LC/MS

Two methods, referred to as Chromatography 1 and Chromatography 2, were alternated using a Thermo Vanquish dual LC coupled to an Orbitrap ID-X. Chromatography 1 used an Acquity BEH Amide analytical column (1.7 μm, 2.1 mm × 150 mm, #176001909, Waters, Eschborn, Germany) combined with a VanGuard pre-column (1.7 μm, 2.1 mm × 5 mm; 186004799, Waters). Mobile phase A consisted of 10% LC/MS grade acetonitrile (A955, Fisher), mobile phase B consisted of 90% LC/MS grade acetonitrile, and both mobile phases contained 10 mM ammonium acetate (73594, Sigma) and 0.2% acetic acid (A11350, Fisher). Column temperature was kept at 40°C, flow rate 0.4 mL/min, and the solvent gradient was as follows: 0-9 min from 95% B to 70% B, 9-13 min from 70% B to 30% B, and 13-14 min held at 30% B followed by a 20 min re-equilibration gradient to prep the column before the next sample injection. The amide column re-equilibration gradient was as follows: 0-1 min held at 30% B at 0.4 mL/min, 1-3 min from 30% B to 65% B and 0.4 mL/min to 0.8 mL/min, 3-15 min held at 65% B and 0.8 mL/min, 15-15.5 min from 65% B to 100% B at 0.8 mL/min, 15.5-17 min held at 100% B at 0.8mL/min, 17-17.5 min from 100% B to 95% B and 0.8 mL/min to 1.2 mL/min, 17.5-19.4 min held at 95% B and 1.2 mL/min, 19.4-19.8 min held at 95% B and decreased from 1.2 mL/min to 0.4 mL/min, and 19.8-20 min held at 95% B and 0.4 mL/min.

Chromatography 2 was a reverse phased chromatography using a CORTECS T3 column (1.6 μm, 2.1mm × 150mm, 186008500, Waters, Eschborn, Germany) combined with a VanGuard pre-column (1.6 μm, 2.1 mm × 5 mm, 186008508, Waters). Mobile phase A consisted of LC/MS grade water (W6-4, Fisher), mobile phase B consisted of 99% LC/MS grade acetonitrile, and both mobile phases contained 0.1% FA (A11710X1-AMP, Fisher). Column temperature was kept at 40°C, flow rate 0.3 mL/min, and the solvent gradient was as follows: 0-10 min from 0% B to 30% B, 10-16 min from 30% B to 100% B, and 16-20 min held at 100% B followed by a 14 min re-equilibration gradient. The T3 column re-equilibration gradient was as follows: 0-6 min held at 100% B and 0.3 mL/min, 6-8 min from 100% B to 0% B at 0.3 mL/min, and 8-14 min held at 0% B and 0.3 mL/min.

For both methods, data were collected with an Orbitrap IDX using an H-ESI source in positive mode. For Chromatography 1, the mass spectrometer parameters were: source voltage 3500V, sheath gas 60, aux gas 19, sweep gas 1, ion transfer tube temperature 300°C, and vaporizer temperature 250°C. Full scan data was collected using the orbitrap with scan range of 105-1000 m/z at a resolution of 120,000. Fragmentation was induced in the orbitrap with assisted HCD collision energies at 20, 40, 60, 80, 100% and with CID collision energy fixed at 35%. For chromatography 2, the mass spectrometer parameters were: source voltage 3500V, sheath gas 70, aux gas 25, sweep gas 1, ion transfer tube temperature 300°C, and vaporizer temperature 250°C. Full scan data was collected using the orbitrap with scan range of 105-1200 m/z at a resolution of 240,000. For both methods, ddMS2 data was acquired with MS1 orbitrap mass resolution at 120,000 and orbitrap MS2 mass resolution at 30,000 for a total cycle time of 0.6 s.

#### Targeted metabolomics data analysis

Peak picking and integration was conducted in Skyline (version 23.1.0.268) using in-house curated compound data bases of accurate mass MS1 and retention time derived from analytical standards and/or MS2 spectral matches on each chromatography method (**Supplemental Table S2**). Full scan raw data files for all samples of a given experiment were imported and metabolite peaks were auto-integrated based off method-specific, in-house curated compound databases that included molecular formula, precursor adducts, and explicit retention times collected from on-method analyzed chemical standards. Manual peak integrations were performed as necessary to account for any tailing peaks, over-lapping peaks, and minor retention time shifting (if any). If manual integration was performed, the “synchronize integration” function was utilized such that all metabolite integration windows were identical between all sample files. Peak areas were then exported for further data processing and analysis. Data were Log-transformed prior to performing statistical analysis. In cases where the same compound is detected on multiple methods, the method with the lowest relative standard deviation in the pooled sample was selected for that compound. Significant differences were determined by ANOVA with a 5% FDR.

#### Untargeted metabolomics data analysis

Compound Discoverer (Thermo Fisher, version 3.3) was utilized for untargeted data analysis. Detected features were assigned putative identifications using MS2 spectral library matching to mzCloud and NIST2020 databases. When no MS2 matches were made, MS1 based chemical formula prediction and accurate mass matches to Chemspider were used for lower-confidence tentative identifications. Features without identification were identified as m/z at retention time (RT). The feature/compound list was refined using blank and peak rating filters. Peak areas for putative compounds passing these filters were subjected to statistical analysis as described above.

### Statistical Analysis

Differential abundance of proteins was assessed using LIMMA eBayes via the R v4.3 (https://cran.r-project.org/) package *limma*. Only proteins without missing values in a given pairwise contrast were included in this analysis and were Log2 transformed. P-values were multiple testing adjusted via Benjamini-Hochberg FDR corrections [68] across all contrasts in a given assay (e.g., corrected across all 3 sets of concentration comparisons AMW20 *versus* AMW35, AMW20 *versus* AMW 50, and AMW35 *versus* AMW50). To assess proteins with missing data, missing values were imputed with the arbitrary low value of one. This imputation was chosen as missing data was assumed to be missing due to being below the limit of detection and thus these proteins would rank lower than observed abundances. These imputed data were then analyzed via semi-parametric ordinal regression from the R package *ordinal* [69]. P-values for these models were calculated via likelihood ratio tests due to the small sample size and were adjusted via Benjamini-Hochberg multiple testing corrections. Gene set enrichment analysis on sets of differentially abundant proteins were conducted via the R package *clusterProfiler* [70]. More specifically, enrichments were based on gene ontology of proteins and focused on biological processes. Significance was based on the ‘optimized FDR q-values’ calculated within *clusterProfiler*. Protein lists were converted to Entrez-ids via org.Mm.eg.db for mice [71] and org.Hs.eg.db for humans [72] (version 3.18.0 for both) All other data were analyzed via Welch’s t-test (2 groups) or F-test (3- or more groups) to account for any inequality of variances.

#### Cheminformatics

Trends in compound hydrophobicity and recovery under modified forms of the AMW extraction procedure were assessed in the non-redundant data subset. 242 of the 252 compounds detected by metabolomics were mapped to human metabolome database (HMDB) or PubChem IDs. Identifiers are unavailable for ten lipids in the dataset and these compounds were excluded. The set of 242 mapped compounds was subsequently filtered to remove redundancy, defined here as cases where the same compound was detected and quantified using two or more methods. This processing step avoids overrepresentation of a subset of chemicals (one-to-many relationships) that would otherwise bias the cheminformatics analysis. A total of 47 of 242 cases of redundancy were identified. These issues were resolved by including data from a single method that yielded the highest quality measurements for a given compound. Specifically, measurements were included for the method that minimized the coefficient of variation (CV) of pooled quality control (QC) samples. These processing steps resulted in a dataset of 195 compounds for understanding compound property-recovery relationships. The HMDB ID or PubChem ID was used to obtain the SMILES (Simplified Molecular Input Line Entry System) via a web-scraping routine implemented using the python request library. The octanol-to-water coefficient (LogP) was predicted from SMILES descriptors using the python RDkit library. For each compound, the Log2 fold change (Log2 FC) of the mean pool size (n = 3) in the experimental condition (AMW25, AMW30, AMW35, AMW40, AMW45, AMW50, AMW55, AMW60) was calculated with respect to the control condition (AMW20). Trends in the Log2 FC and the LogP were analyzed by computing Spearman’s rank correlation coefficient (non-parametric. Correlation coefficients were calculated using the python scipy.stats module. All cheminformatics scripts were developed using python v3.9.7.

## Acknowledgments

We would like to thank Drs. Russell Jones, Sara Nowinski, Evan Lien, and Nick Burton for their critical evaluation of this work. We also thank Megan Gendjar and Lisa DeCamp for their experimental assistance, and Drs. Matt Steensma and Carrie Graveel for providing HEK cells and tissue culture reagents.

## Author contributions

Conceptualization: RJH, MTSH, RDS

Methodology: RJH, MTSH, CDC, ZBM, AEE, CNI, CDC, ABJ, MPV, HL, RDS

Formal Analysis: RJH, MTSH, CDC, ZBM, EW, MPV, RDS

Investigation: RJH, MTSH, CDC, AEE, CNI, ABJ, MPV, RDS

Resources: HL, RDS

Visualization: RJH, MTSH, ZBM, EW, MPV, KW

Supervision: RDS

Project Administration: MLEG, KW, RDS

Funding acquisition: RDS

Writing—original draft: RJH, MTSH, MLEG, RDS

Writing—review & editing: RJH, MTSH, CDC, ZBM, EW, MPV, AEE, CNI, CDC, ABJ, MLEG, KW, HL, RDS

## Data and materials availability

Proteomics data have been deposited on ProteomeExchange (https://www.proteomeexchange.org). Metabolomics data have been included as supplemental files, with raw MS data files made available on request. All other data are available in the main text or the supplementary materials.

