## Supplementary Figures and Table 5 for "A diverse proteome is present and enzymatically active in metabolite extracts"

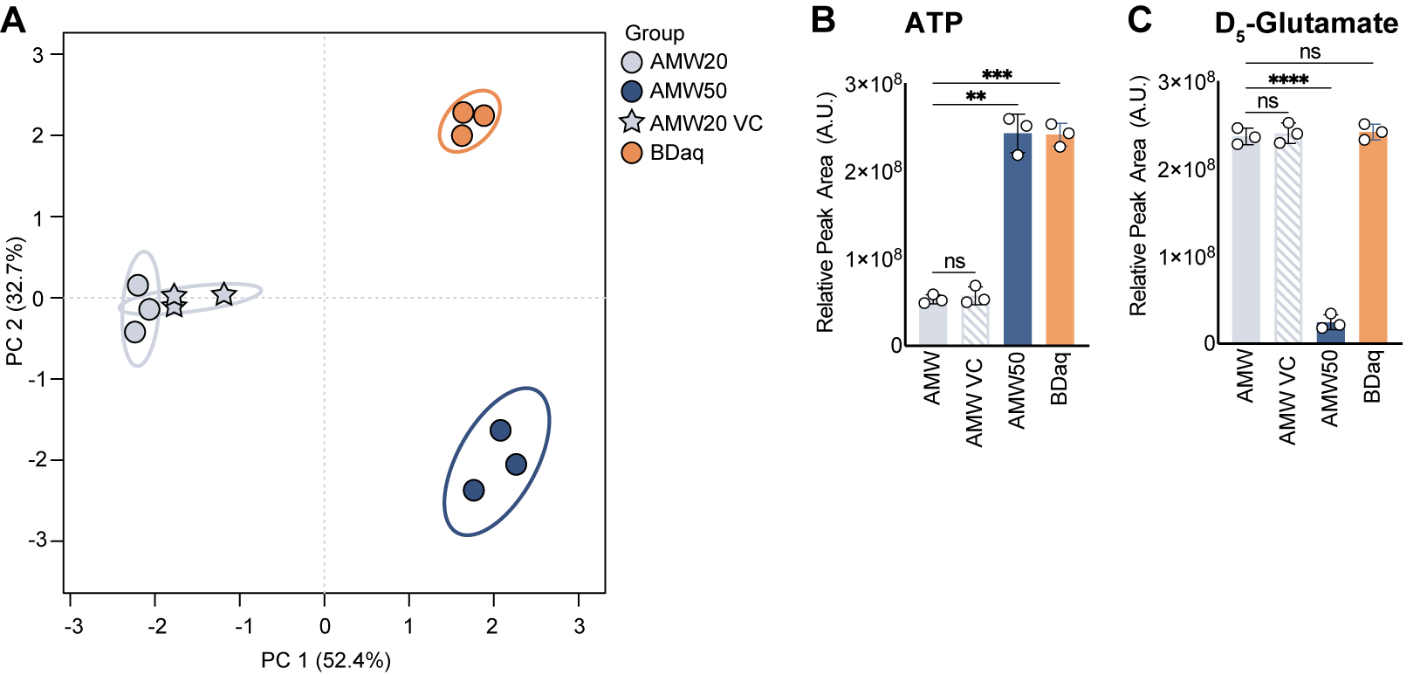

Fig. S1.

Extraction water content impacts metabolite extraction independent of extraction volume. (A) PCA of murine liver metabolites across AMW20, AMW20 VC (final volume equal to that of AMW50 condition, but secondary incubation was with the AMW20 solvent mixture instead of H_2_O), AMW50, and Bligh Dyer aqueous (BDaq) extraction conditions (MetaboAnalyst 6.0, 95% confidence ellipse, *n*=3 per group). (B) Relative abundance of ATP in murine liver across AMW20, AMW20 VC, AMW50, and BD extraction conditions. Significance calculated by Welch ANOVA (mean ± SD, *n*=3 per group). (C) Relative abundance of D_5_-glutamate in murine liver across AMW20, AMW20 VC, AMW50, and BD extraction conditions. D_5_-glutamate was added at resuspension. Significance calculated by Welch ANOVA (mean ± SD, *n*=3 per group).

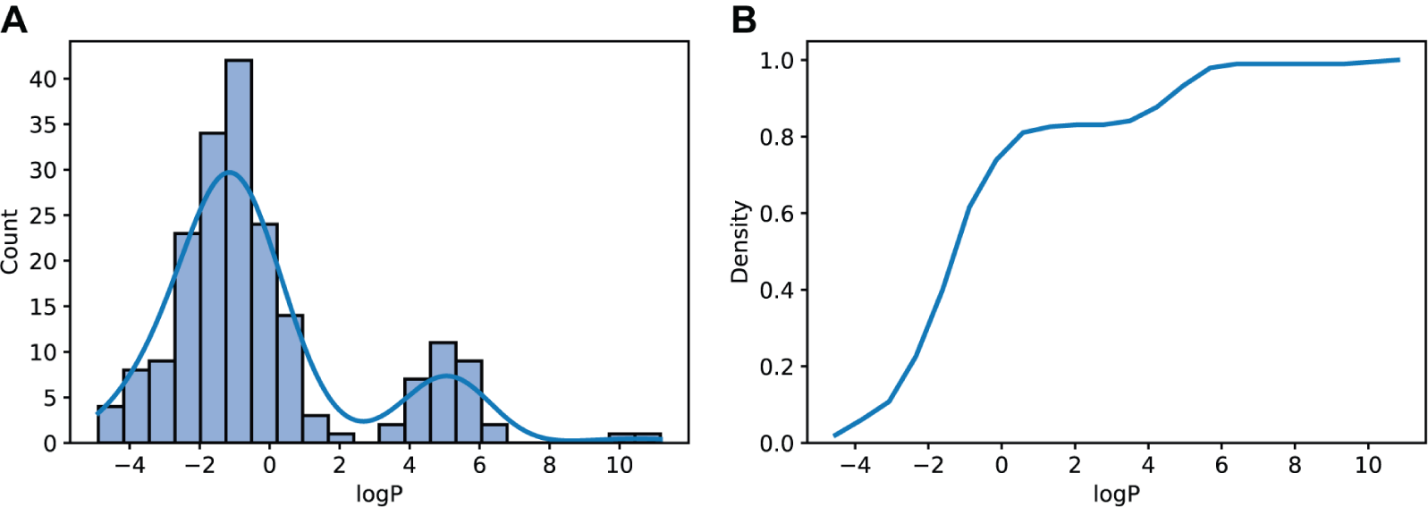

Fig. S2.

Metabolite LogP distribution. (A) Histogram of the octanol-water coefficient (LogP) distribution in the cheminformatics analysis. The kernel density estimate (KDE) is overlayed. (B) The cumulative distribution function is shown.

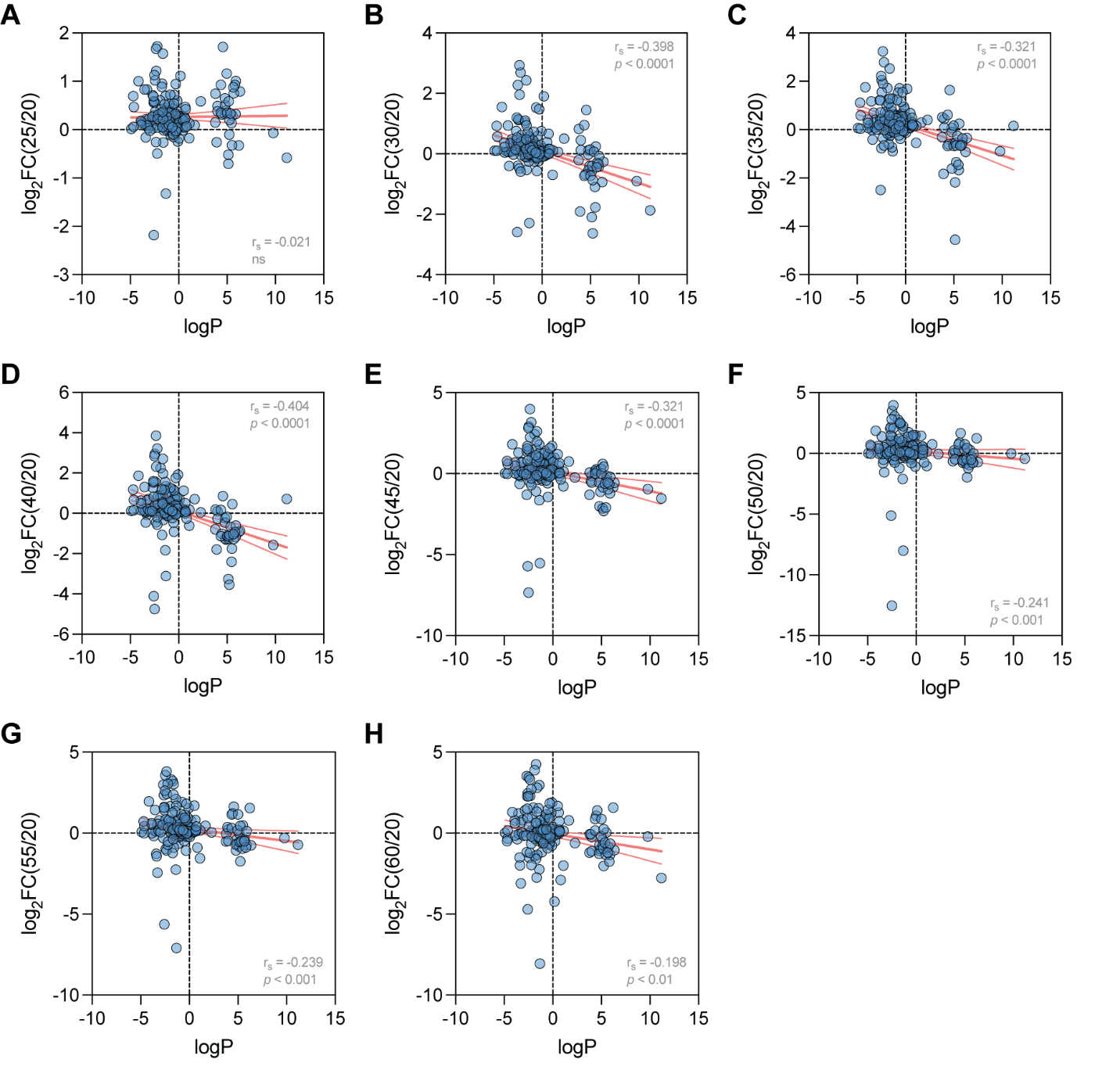

Fig. S3.

**Correlation between metabolite logP and recovery under the specified extraction. conditions.** The mean Log2 fold change (*n*=3) of metabolite levels in the (**A**) AMW25, (**B**) AMW30, (**C**) AMW35, (**D**) AMW40, (**E**) AMW45, (**F**) AMW50, (**G**) AMW55, (**H**) AMW60 extraction condition is shown with respect to the control condition (AMW20). In all cases, the Spearman correlation coefficient (r_s_) was calculated for the specified Log2 fold change and LogP datasets. These values are inset in each plot together with the p-value, simple linear regression models (red solid lines) and the 95% confidence intervals (red dashed lines) are displayed. Abbreviations: ns = not significant (determined using a 5% significance level).

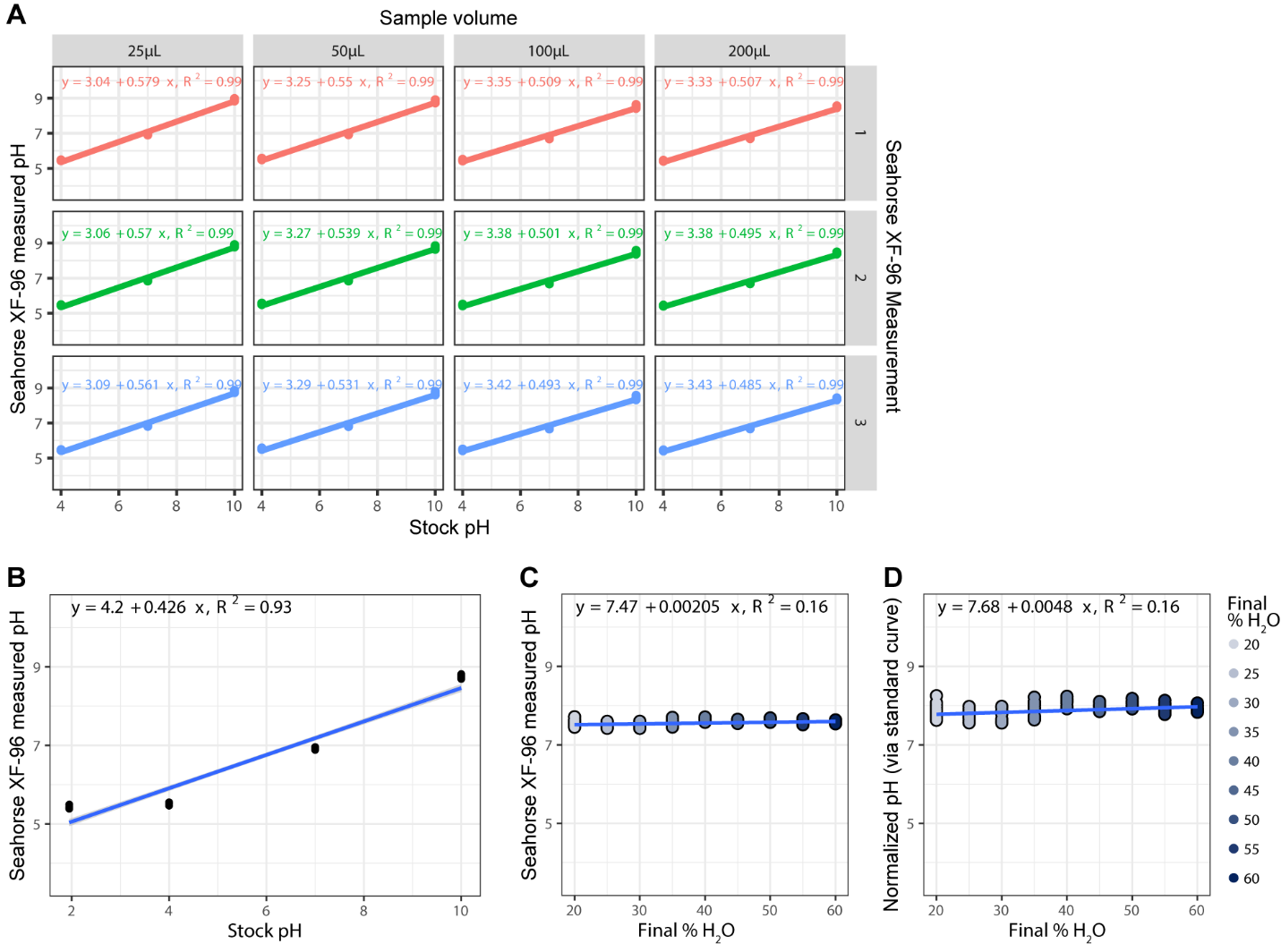

Fig. S4.

**Extraction water content impacts metabolite extraction independent of pH.** (**A**) The Seahorse XF-96 is an accurate micro-pH meter for 25-200 μL volumes. Linear regression models (solid lines) and equations are displayed across measurements (1, 2, and 3) and sample volumes (25, 50, 100, and 200 μL). The *R^2^* value comparing stock pH and Seahorse F-96-measured pH values are shown. (**B**) Standard curve for the AMW20-AMW60 pH analysis. Linear regression model (solid lines) and equation are displayed. The *R^2^* value comparing stock pH and Seahorse F-96-measured pH values are shown. (**C**) Extraction water content (AMW20-AM60) and raw resuspension pH values do not correlate. Linear regression model (solid lines) and equation are displayed. The *R^2^* value comparing stock pH and Seahorse F-96-measured pH values are shown. (**D**) Extraction water content (AMW20-AM60) and standard-curve-normalized resuspension pH values do not correlate. Linear regression model (solid lines) and equation are displayed. The *R^2^* value comparing stock pH and Seahorse F-96-measured pH values are shown.

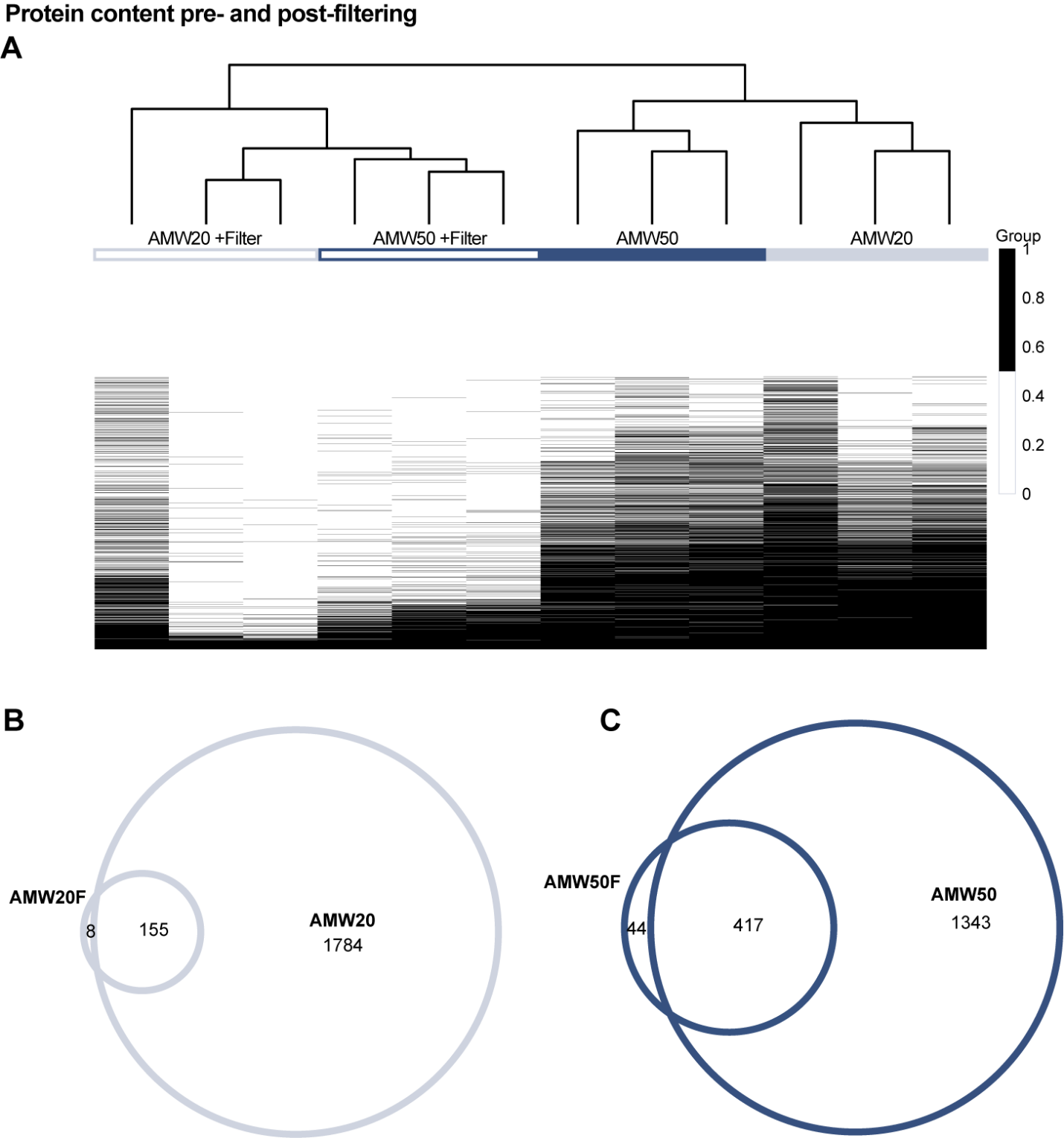

Fig. S5.

**3 kDa filtration removes proteins from metabolite extracts.** (**A**) Heatmap depicting relative abundance of proteins across AMW20, AMW20 +Filter (AMW20F), AMW50, and AMW50 +Filter (AMW50F) extraction conditions (**B**) Venn diagram of AMW20 and AMW20F overlapping proteins. (**C**) Venn diagram of AMW50 and AMW50F overlapping proteins.

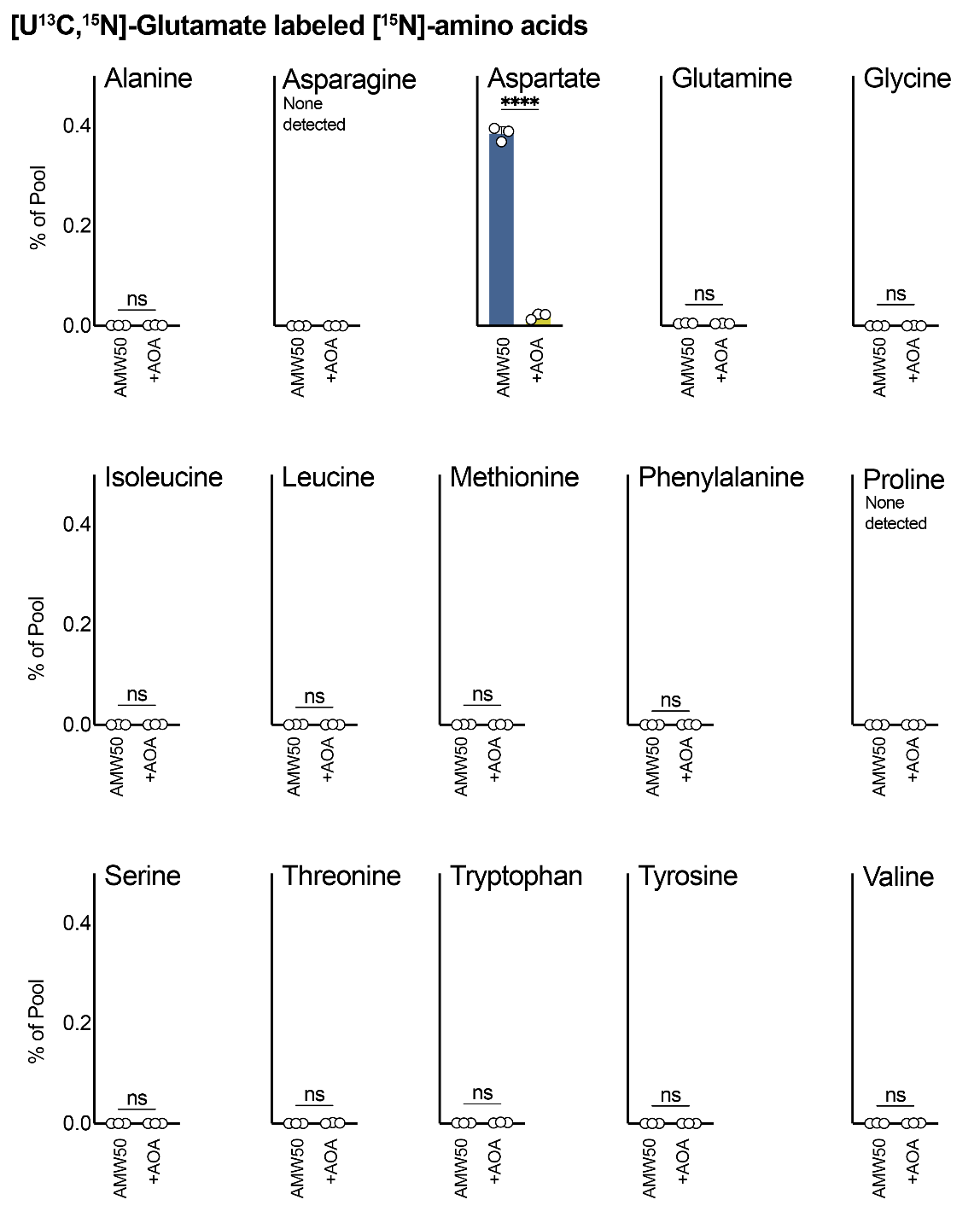

Fig. S6.

**[U^13^C,^15^N]-glutamate incorporation into amino acids.** Relative abundance of [U^13^C,^15^N]-glutamate -derived amino acids in murine liver in AMW50 and AMW50 +AOA extraction conditions. Significance calculated by Welch t-test (mean ± SD, *n*=3 per group).

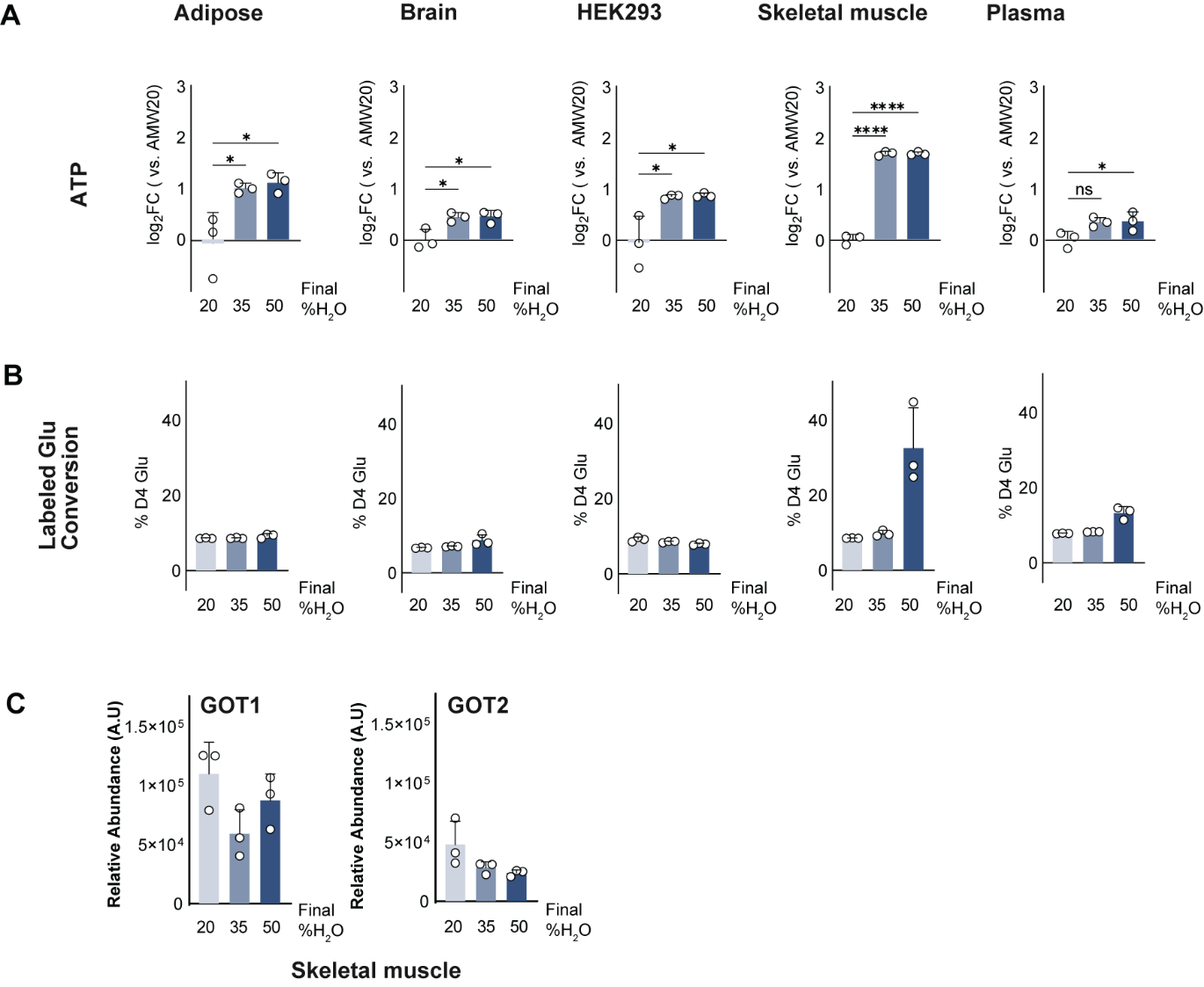

Fig. S7.

**Multi-matrix assessment of ATP, D_5_ Glu to D_4_ Glu conversion, and GOT1/GOT2.** (**A**) Log2 fold change in ATP signal at 3 final water concentrations *versus* the average of AMW20 for adipose, brain, HEK293 cells, skeletal muscle, and plasma matrices. Significance calculated by Welch ANOVA (mean ± SD, *n*=3 per group). (**B**) Percentage of D_4_-glutamate as a portion of the total labeled glutamate pool (D_4_ glu + D_5_ glu signal) for adipose, brain, HEK293 cells, skeletal muscle, and plasma matrices in AMW20, AMW35 and AMW50 extracts (mean ± SD, *n*= 3 per group). (**C**) Relative abundance of GOT1 and GOT2 proteins in skeletal muscle as a function of water content (mean ± SD, *n*= 3 per group).

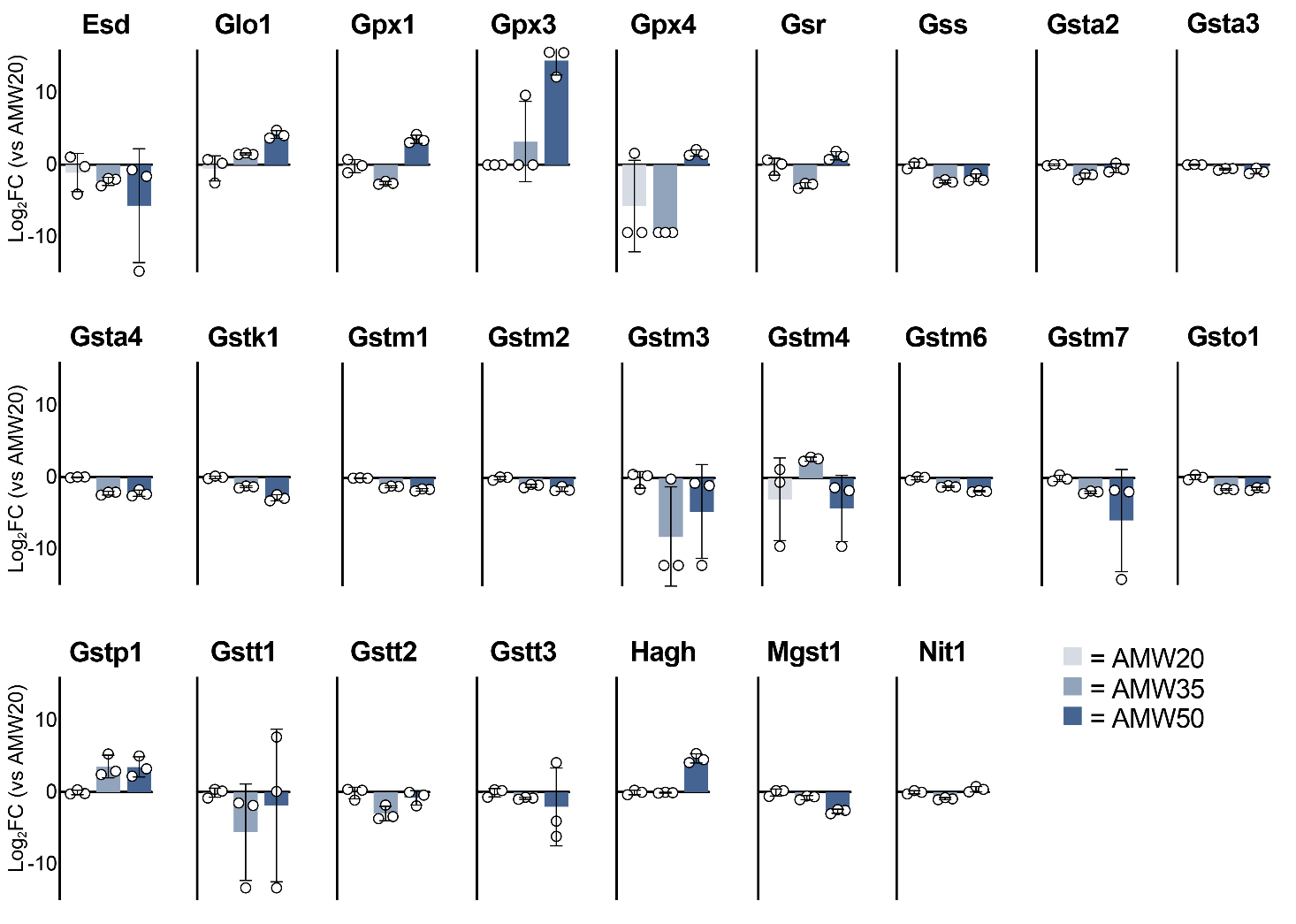

Fig. S8.

**Glutathione-related proteins are present in murine liver metabolite extracts.** (**A**) Relative abundance of murine liver glutathione-related proteins in AMW20, AMW35, and AMW50 samples (mean ± SD, *n*=3 per group).

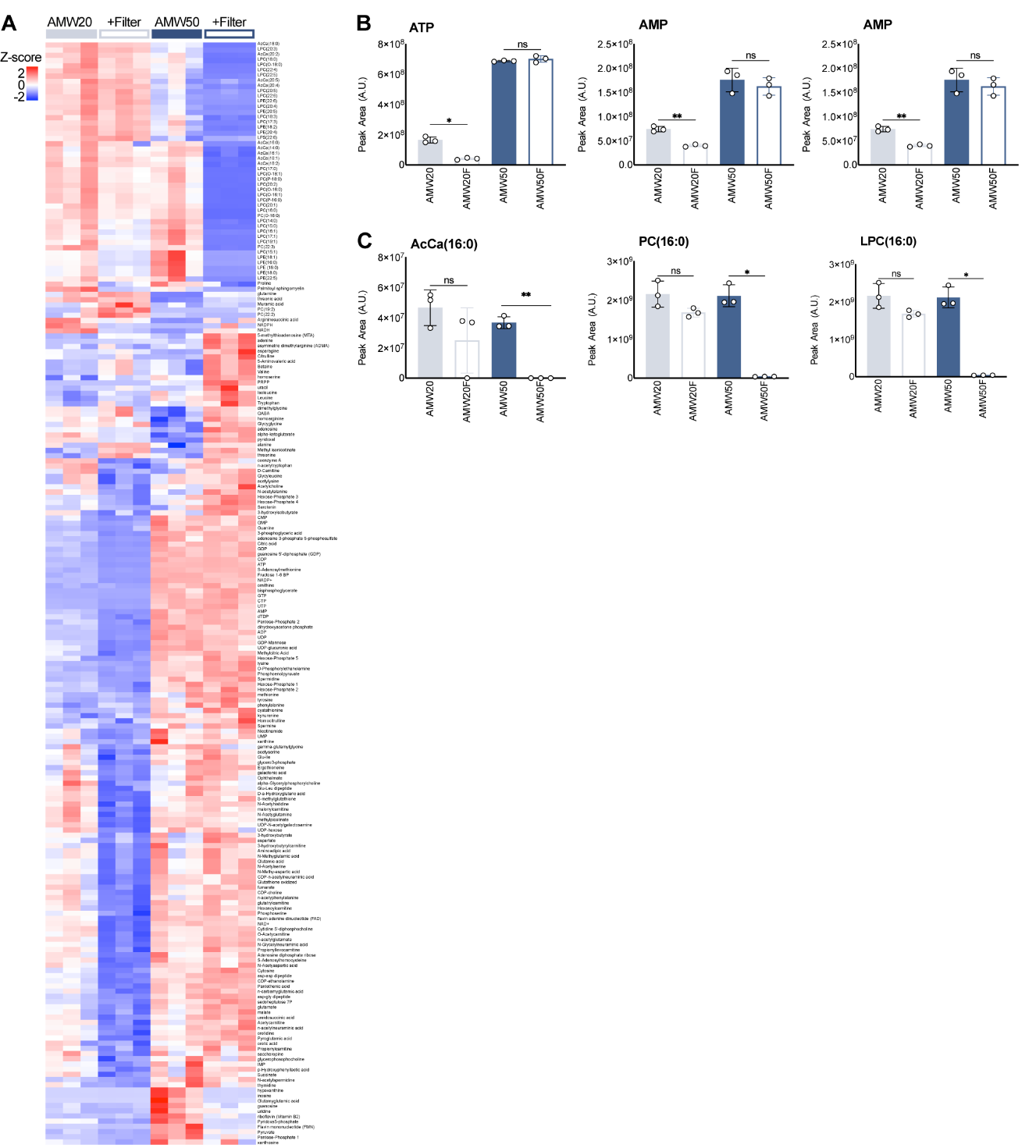

Fig. S9.

**Metabolic effects of filtration in AMW20 and AMW50 conditions.** (A) Heatmap depicting relative abundance of murine liver metabolites across AMW20, AMW20 +Filter (AMW20F), AMW50, and AMW50 +Filter (AMW50F) extraction conditions. All significantly different metabolites are shown (**Figure 9A**) and labeled. Significance was calculated by ANOVA using MetaboAnalyst 5.0, and row hierarchical clustering was performed using Morpheus (Broad). (**B**) Relative abundance of ATP, ADP, and AMP in murine liver across AMW20, AMW20F, AMW50, and AMW50F extraction conditions. Significance calculated by Welch ANOVA (mean ± SD, *n*=3 per group).  **(C)** Relative abundance of AcCa(16:0), PC(16:0), LPC(16:0) in murine liver across AMW20, AMW20F, AMW50, and AMW50F extraction conditions. Significance calculated by Welch ANOVA (mean ± SD, *n*=3 per group).

The following tables have been uploaded as .xlsx files

Table S1.

Metabolomics extraction worksheet

Table S2.

Metabolomics compound list

Table S3.

Metabolite profile of AMW20, AMW35 and AMW50 from adipose, brain, skeletal muscle, HEK293 cells and mouse plasma

Table S4.

Proteomics profile of AMW20, AMW35 and AMW50 from adipose, brain, skeletal muscle, HEK293 cells and mouse plasma

| **REAGENT or RESOURCE** | **SOURCE** | **IDENTIFIER** |
| --- | --- | --- |
| **Chemicals and reagents** | | |
| 1X PBS | Gibco | 10010023 |
| Acetic Acid, Optima™ LC/MS | Fisher Chemical | A11350 |
| Acetonitrile, Optima™ LC/MS Grade | Fisher Chemical | A955-4 |
| Aminooxyacetic Acid | Cayman Chemical | 28298 |
| Ammonium acetate (LiChropur™) | Millipore Sigma | 73594-100G-F |
| Chloroform, LiChrosolv | Millipore Sigma | 1024441000 |
| DMEM, no phenol red | Gibco | 21063029 |
| EDTA | VARI Media Lab | 100295 |
| Fetal bovine serum | VWR | 1300-500 |
| Formic Acid, 99.0+%, Optima™ LC/MS Grade | Fisher Chemical | A11710X1-AMP |
| InfinityLab deactivator additive (Medronic acid) | Agilent Technologies | 5191-4506 |
| L-Glutamic acid (¹³C₅, 99%; ¹⁵N, 99%) | Cambridge Isotope Laboratories | CNLM-554-H-0.25 |
| L-Glutamic Acid (2,3,3,4,4-D_5_, 97-98%) | Cambridge Isotope Laboratories | DLM-556 |
| L-Tryptophan (D_8_, 97-98%) | Cambridge Isotope Laboratories | DLM-6903-0.25 |
| Methanol, Optima™ LC/MS Grade | Fisher Chemical | A456-4 |
| NaCl | Gibco | S8776 |
| Penicillin-streptomycin | Gibco | 15070063 |
| Tributylamine | Millipore Sigma | 90780-100ML |
| Water with 0.1% formic acid, Optima™ LC/MS Grade | Fisher Chemical (via Block Scientific) | LS118-1 |
| Water with 0.1% TFA, Optima™ LC/MS Grade | Fisher Chemical (via Block Scientific) | LS119-500 |
| Water, Optima™ LC/MS Grade | Fisher Chemical | W6-4 |
| **Supplies** | | |
| 1.5 mL Eppendorf tubes | DOT Scientific | RN1700-GMT |
| 1.5 mL Protein LoBind® Tubes | Eppendorf | 30108442 |
| 1.5 mL screw cap micro tube, low protein binding | SARSTEDT | 72.703.600 |
| 2.0 mL Protein LoBind® Tubes | Eppendorf | 30108450 |
| 5 mL Eppendorf tubes | Eppendorf | 30119401 |
| 6-well tissue culture plates | Corning | 3506 |
| 96 well Clear Flat Bottom Plates | Costar (Corning) | 3370 |
| 96 Well V Bottom Plates | Greiner | 651201 |
| Adhesive 96 well plate seals | Bio-Rad | MSB1001 |
| Amicon Ultra-2 3K centrifugal filter devices | Millipore Sigma | UFC200324 |
| Cell scrapers | Fisher | 08-100-240 |
| Ceramic homogenizer tubes | OMNI International Inc. | 19-627 |
| Certified QSertVial™ (vial with fused-in insert) | Millipore Sigma | 29391-U |
| Certified Vial Kit, Low Adsorption (LA), 2 mL | Millipore Sigma | 29651-U |
| epT.I.P.S.® 0.5-20 μL | Eppendorf | 22492021 |
| epT.I.P.S.® 2-200 μL | Eppendorf | 22492039 |
| epT.I.P.S.® 20-300 μL | Eppendorf | 22492047 |
| epT.I.P.S.® 50-1000 μL | Eppendorf | 22492055 |
| LC autosampler caps |  | 6PSC9STS1R |
| LC autosampler vials |  | 6PSV9-03FIVP |
| **Critical commercial assays** | | |
| EasyPep Mini MS Sample Prep Kit | Thermo Scientific | A40006 |
| Pierce BCA Protein Assay Kit | Thermo Scientific | 23225 |
| **Experimental models (cell lines)** | | |
| Phoenix-AMPHO | ATCC | CRL-3213 |
| **Experimental models (organisms/strains)** | | |
| Mouse: C57BL/6J | The Jackson Laboratory | RRID: IMSR_JAX:000664 |
| **Software and algorithms** | | |
| Compound Discoverer 3.3 SP2 | Thermo Fisher Scientific | OPTON-31061 |
| Graphpad Prism | GraphPad Software | [https://www.graphpad.com](https://www.graphpad.com/) |
| MetaboAnalyst 5.0 and 6.0 | MetaboAnalyst | <https://www.metaboanalyst.ca/> |
| R studio | Posit | <https://www.rstudio.com/products/rstudio/> |
| Skyline 23.1 | MacCoss Lab Software | <https://skyline.ms/project/home/begin.view> |
| Spectronaut v18 | Biognosys | <https://biognosys.com/resources/spectronaut-18/> |

Table S5.

**List of reagents and resources used in this study.** Including manufacturer and catalogue number.
